# Cryo-EM reveals open and closed Asgard chromatin assemblies

**DOI:** 10.1101/2025.05.24.653377

**Authors:** Harsh M. Ranawat, Marc K.M. Cajili, Natalia Lopez-Barbosa, Thomas Quail, Remus T. Dame, Svetlana O. Dodonova

## Abstract

Asgards are the closest archaeal relatives of eukaryotes, representing an important step in chromatin evolution. However, their chromatin organization has remained enigmatic until now. In this study, we present the first structures of Asgard chromatin assemblies formed by the Hodarchaeal histone HHoB. Our high-resolution cryo-EM structures reveal that this Asgard histone assembles into compact "closed" and into extended "open" hypernucleosomes. Thus the closed hypernucleosome conformation is conserved across archaeal lineages, while the open conformation resembles a eukaryotic H3-H4 octasome and likely represents an Asgard-specific innovation. Moreover, we show that Mg²⁺ ions influence Asgard chromatin conformation, suggesting a regulatory role. Overall, our study provides the first structure-based model of Asgard chromatin organization, expanding our understanding of chromatin architecture in evolutionary context.

## Introduction

Eukaryotes and most Archaea use histone proteins to organize their genomes and regulate chromatin states. In eukaryotes, a histone octamer wraps 147 bp of DNA into a nucleosome^1^. The histone fold domain and dimer handshake motif are conserved across the tree of life^2,3^, and it is widely accepted that the eukaryotic histones trace their lineage to archaeal histones^4^. However, while eukaryotic core histones form obligate heterodimers, archaeal histones can form both homo- and heterodimers^5^. Furthermore, instead of forming nucleosomes of a defined size (∼147 bp), archaeal histones assemble into histone-DNA complexes “nucleosomes” of variable size (N × 30 bp), where N histone dimers can each wrap 30 bp of DNA^6,7^. Notably, most archaeal histones lack the extended tails that are hallmarks of eukaryotic histones^8^.

Our current understanding of archaeal chromatin is largely based on a limited number of model species, primarily from the Euryarchaeota branch, such as *Methanothermus fervidus* and *Thermococcus kodakarensis*. The first pivotal study on archaeal chromatin structure revealed a 90 bp archaeal nucleosome formed by “classical” archaeal tail-less histone HMfB from *M. fervidus*^6^. Hereafter, we refer to any archaeal histone-DNA complex as a “nucleosome” regardless of the size or number of histones bound, and we refer to nucleosomes with multiple superhelical turns (more than 2) – as “hypernucleosomes”. Interestingly, the crystal packing of this structure displayed a continuous, tightly packed "hypernucleosome" with strong stacking interactions between histones^8^, which were also later demonstrated via biophysical approaches^9^. Subsequently, a cryo-electron microscopy (cryo-EM) study of a similar HTkA histone from *T. kodakarensis* confirmed this structure and revealed an additional "slinkie-like" configuration^10^. In the "slinkie" structure, a 90 bp nucleosome was connected via a short DNA linker to a 120 bp nucleosome, while the arrangement of histones within these nucleosomes remained consistent with the HMfB-based structure. This study demonstrated that archaeal chromatin exhibits greater conformational diversity than previously recognized.

Although Euryarchaeota provided early insights into archaeal chromatin, many other archaeal groups remain understudied, which limits our comprehensive understanding of archaeal chromatin architecture. The recently discovered Asgard superphylum of archaea presents particularly intriguing chromatin features based on metagenomic analyses^11^. Many Asgard metagenomes encode multiple histone variants, some of which include extended histone tails^8^. For instance, the Asgard metagenome of Hodarchaeales LC3 (previously classified as *Candidatus Heimdallarchaeota LC3*^12^) encodes ten histone variants: eight short tail-less histones and two histones with extended tails^8,11^. Asgard archaea typically possess larger genomes and encode eukaryotic signature proteins^12^. It is widely accepted that eukaryotes originated from within the Asgard superphylum^12–14^, inheriting much of their "information processing" machinery from archaea. Consequently, Asgard histones represent a critical step in chromatin evolution, leading toward eukaryotic chromatin or to viral histones as an intermediate step^11,15,16^.

In this study, we provide the first insights into the structure and function of Asgard chromatin assemblies. We structurally, biochemically, and biophysically characterize chromatin assemblies formed by a tail-less histone (HHoB) from the Asgard archaea Hodarchaeales LC3. We present several high-resolution cryo-EM structures of nucleosomes formed by the HHoB histone wrapping DNA. Our structures reveal that the HHoB histone forms two distinct nucleosomal conformations - "closed" and "open" - under a wide range of conditions. We demonstrate that increasing Mg²⁺ concentrations favor the formation of continuous hypernucleosomes, and we report cryo-EM structures of both closed and open hypernucleosomes. The ability of Asgard histones to form closed nucleosomes and hypernucleosomes experimentally confirms that this chromatin state is conserved across distant archaeal groups such as Euryarchaeota and Asgards. In contrast, the novel open chromatin state has not been observed previously and may represent an Asgard-specific innovation. Overall, our study provides the first structural insights into Asgard chromatin organization, expanding our understanding of archaeal chromatin architecture. These findings are validated through mutagenesis, biochemical assays, and single molecule biophysical experiments.

## Results

### Asgard histone HHoB forms open and closed nucleosomes in vitro

The LC3 Asgard genome encodes ten histone proteins, the majority of which (eight) are short histones lacking extended tails (Figure 1A). Firstly, three of the short histones (designated HHoB, HHoF, and HHoG) were selected for initial biochemical analysis (Figure 1A). Each histone was recombinantly expressed in *E. coli* and purified via affinity and ion-exchange chromatography. Archaeal nucleosomes were then reconstituted *in vitro* by incubating Widom601 147 bp DNA with each histone in buffer A (20 mM HEPES pH 7.5, 100 mM NaCl). Widom601^17^ DNA was used to allow direct comparison with other structures such as the archaeal HTkA-DNA complexes, which were also reconstituted using Widom601^10^. All three histones exhibited similar DNA binding behavior in electrophoretic mobility shift assays (EMSA) and demonstrated a comparable range of binding affinities (Figure 1B, Figure S1). For HHoB, the affinity was determined to be 30.8 ± 3.7 nM (mean ± standard deviation, N=3) as measured by microscale thermophoresis (MST) (Figure S2). Unlike the well-studied archaeal histone HMfB from the Euryarchaeon *M. fervidus*, which displays highly cooperative binding behavior reflected in a single shifted band^18^, the HHoB, HHoF, and HHoG histones exhibited a ladder-like step-wise binding pattern (Figure 1B, and Figure S1). Based on the similar behavior of these three tail-less Asgard histones, we proceeded with a more in-depth analysis of the HHoB histone.

**Figure 1.**
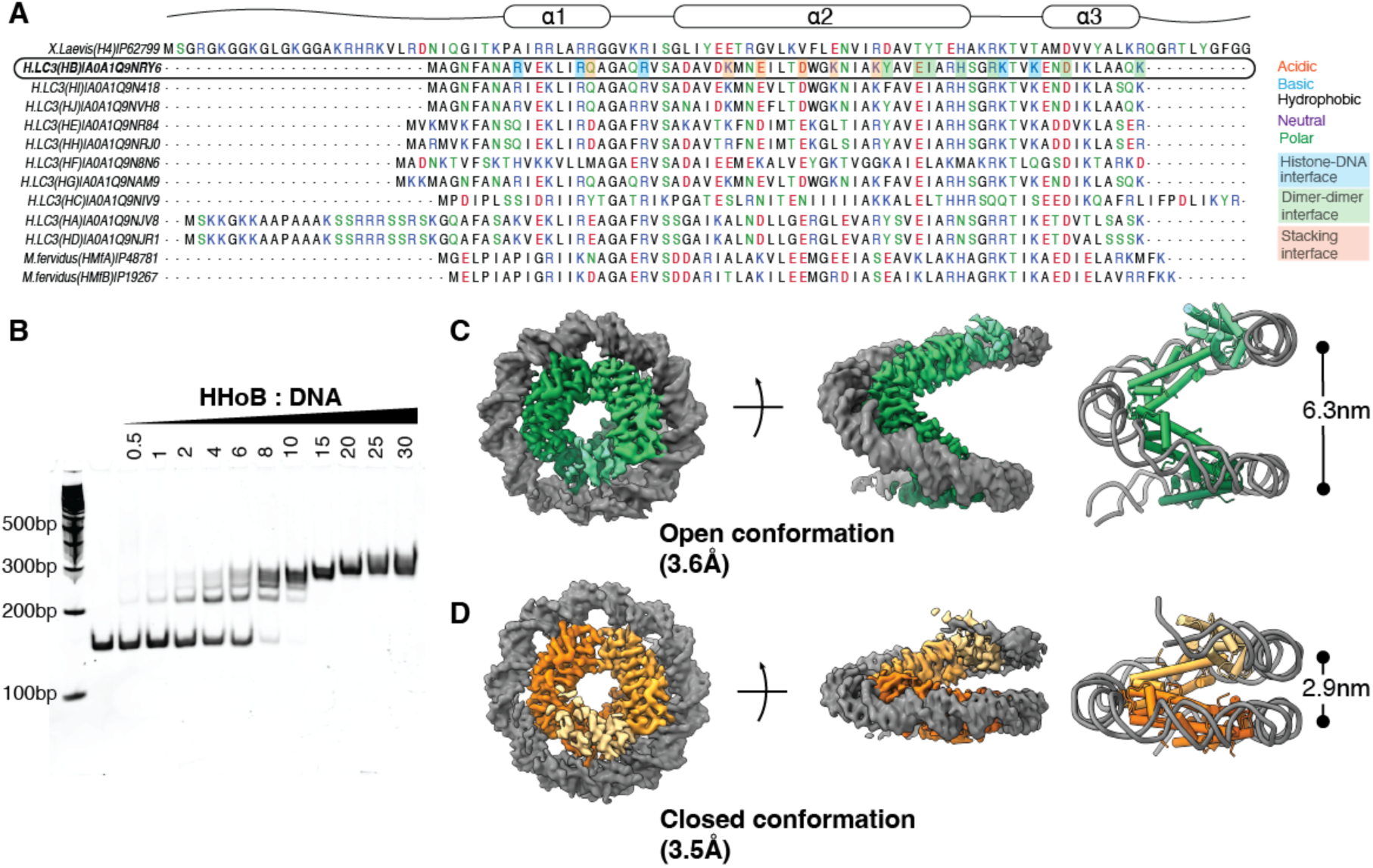
(A) Multiple sequence alignment (MSA) of histones from LC3 Asgard metagenome, the HMfB histone from *M.fervidus*, and histone H4 from *X.laevis*. Histone fold arrangement is shown schematically above. Residues involved in histone-DNA (grey), dimer-dimer (green), stacking interfaces (orange) based on the HHoB-DNA structures are highlighted with colored bars. (B) EMSA gel of Widom601 147 bp DNA with increasing molar ratios of HHoB : DNA. (C, D) HHoB-DNA complex in 1 mM MgCl₂, shown in open (C) and closed (D) conformations. Top and side views of the EM map (left) and molecular model (right) are presented, with DNA in grey and histone dimers in shades of green (C, PDB 9QV7) and orange (D, PDB 9QV5). Superhelical pitch (distance between DNA gyres) is indicated with a bar.

Mg²⁺ ions are essential and abundant in eukaryotic cells (1-10 mM^19,20^), and in archaeal cells (i.e. up to 120 mM in *Thermococcus* Euryarchaea^21^), and are known to modulate chromatin structure in bacteria^22,23^. However, as Asgard archaea are extremely difficult to isolate and cultivate^24,25^, we lack the data on their intracellular ion concentration, and can only assume that the physiological range is anywhere between the known ranges for archaea, bacteria, and eukaryotes (1-120 mM). Notably, samples used in all structural studies in the field have also contained Mg²⁺ ^6,10^. Therefore, we reconstituted the HHoB-based Asgard nucleosomes in buffer A supplemented with 1 mM Mg²⁺, where the concentration of 1 mM was chosen as a minimal physiologically relevant. The samples were incubated at room temperature (RT) for 20 minutes, plunge-frozen, and subjected to cryo-EM single particle analysis (SPA). From the collected particles, after 2D and 3D classification in cryoSPARC^26^, two main 3D classes emerged, revealing two distinct conformations that we refer to as "closed" and "open" (Figure 1C and 1D). Both classes were equally abundant in the dataset (34.7% and 42.4%) along with some ‘low quality’ classes that were filtered out (Figure S3). The two main classes were refined to 3.4 Å and 3.6 Å resolution respectively (Figure S3, Table 1, Video S1), and the corresponding molecular models were built using Phenix^27^, Coot^28^, and ISOLDE^29^ (see Methods).

**Table 1:**
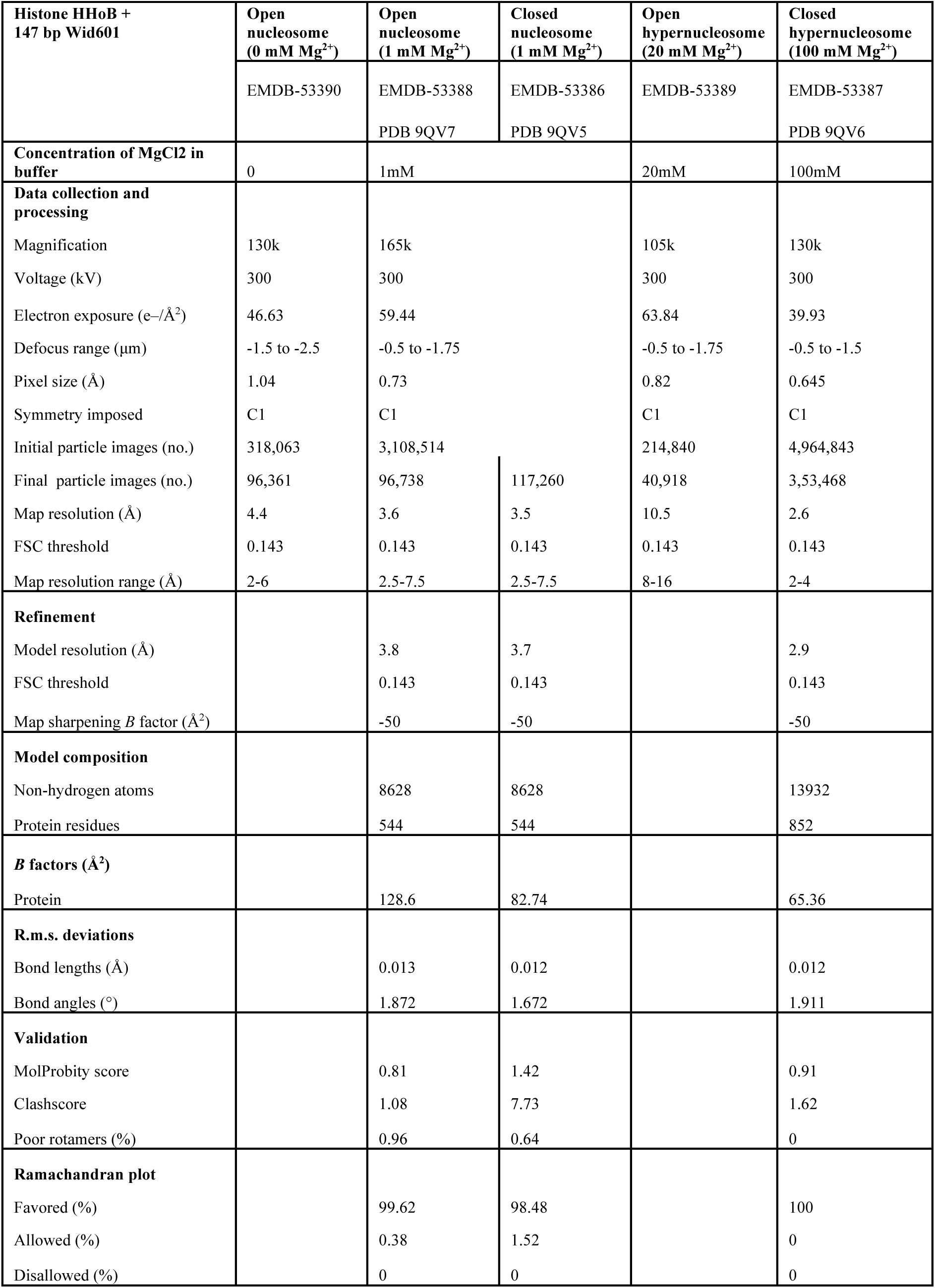
Cryo-EM data collection, refinement and validation statistics.

Each of the cryo-EM maps reveals a well-resolved left-handed nucleosome superhelix (Figure 1C-D) with four HHoB histone dimers bound to 120 bp of the DNA (out of 147 bp total), consistent with the expected 30 bp footprint of a histone dimer. We note that in both EM maps, at higher density visualization thresholds, we also observe the fifth HHoB dimer bound to the DNA, albeit at lower local resolution most likely caused by higher flexibility of the DNA end, which is typical for dynamic nucleosomal complexes^30^ (Figure S3A). Binding of five histone dimers corresponds well to the number of shifts observed in the EMSA assays – where binding of each dimer leads to appearance of a shifted band (Figure 1B). Binding of five histone dimers also shows that the Widom601 sequence does not have any positioning effect on the HHoB, and it’s positioning is simply defined by the length of the available DNA. The footprint of a histone dimer was further verified via EMSA with DNA templates of different lengths (Figure S2B).

The "closed" nucleosome conformation (PDB 9QV5) is similar to the previously reported HMfB euryarchaeal nucleosome^6^ (RMSD 1.09 Å between Cα atoms), while the "open" conformation (PDB 9QV7) is novel and significantly less compacted than the closed conformation (Figure 1C-D). The pitch of the closed nucleosome is 29.5 Å (measured between DNA atoms N1 of A30 and N3 of T105), whereas the pitch of the open nucleosome is almost twice as wide at 63.0 Å (measured between DNA atoms N1 of A30 and N3 of A103).

### Open and Closed HHoB Nucleosomes Co-exist in a Range of Mg²⁺ Conditions and Form Hypernucleosomes

We investigated the influence of increasing Mg²⁺ concentrations on Asgard nucleosome structure, as Mg²⁺ plays a crucial role in chromatin structure and dynamics regulation in both archaea and eukaryotes^31,32^. Based on reported physiological ranges of intracellular Mg²⁺ concentration in archaea^21^, we reconstituted Asgard HHoB nucleosomes in the presence of 0 to 100 mM Mg²⁺ in buffer A (Methods). Each sample was incubated at RT for 20 min, plunge-frozen, and examined by cryo-EM (Figure S4). For each Mg²⁺ concentration (0, 1, 20, 40, 60, 80, 100 mM), at least 30 micrographs were collected. First, we chose the 40 mM dataset that contained all types of particles – open, closed, and mixed (combination of stacking open and closed particles) (Figure 2A). Next, after preprocessing, we trained a Topaz^33^ model to pick particles in the dataset. This model was then used to pick particles from all datasets (with different Mg conditions). Next, “low quality” particles (damaged particles, ice contamination etc.) were removed by 2D classification and sorting. We visualized and calculated the number of open, closed, or mixed 2D classes based on the helical pitch observed in each 2D class (Figure 2). Both closed and open nucleosomes were observed across a wide range of Mg²⁺ concentrations (1-60 mM) (Figure 2A). At higher Mg²⁺ concentrations (80 and 100 mM), only closed nucleosomes were observed (Figure 2), while in the absence of Mg²⁺, only open nucleosomes were detected. We never observed chromatin aggregation or precipitation even at the highest tested Mg²⁺ concentrations (100 mM), whereas eukaryotic chromatin is known to precipitate and aggregate at very low Mg²⁺ concentrations (5 mM)^10^. This property may be crucial for archaea that can have up to 120 mM intracellular Mg²⁺ concentrations.

**Figure 2.**
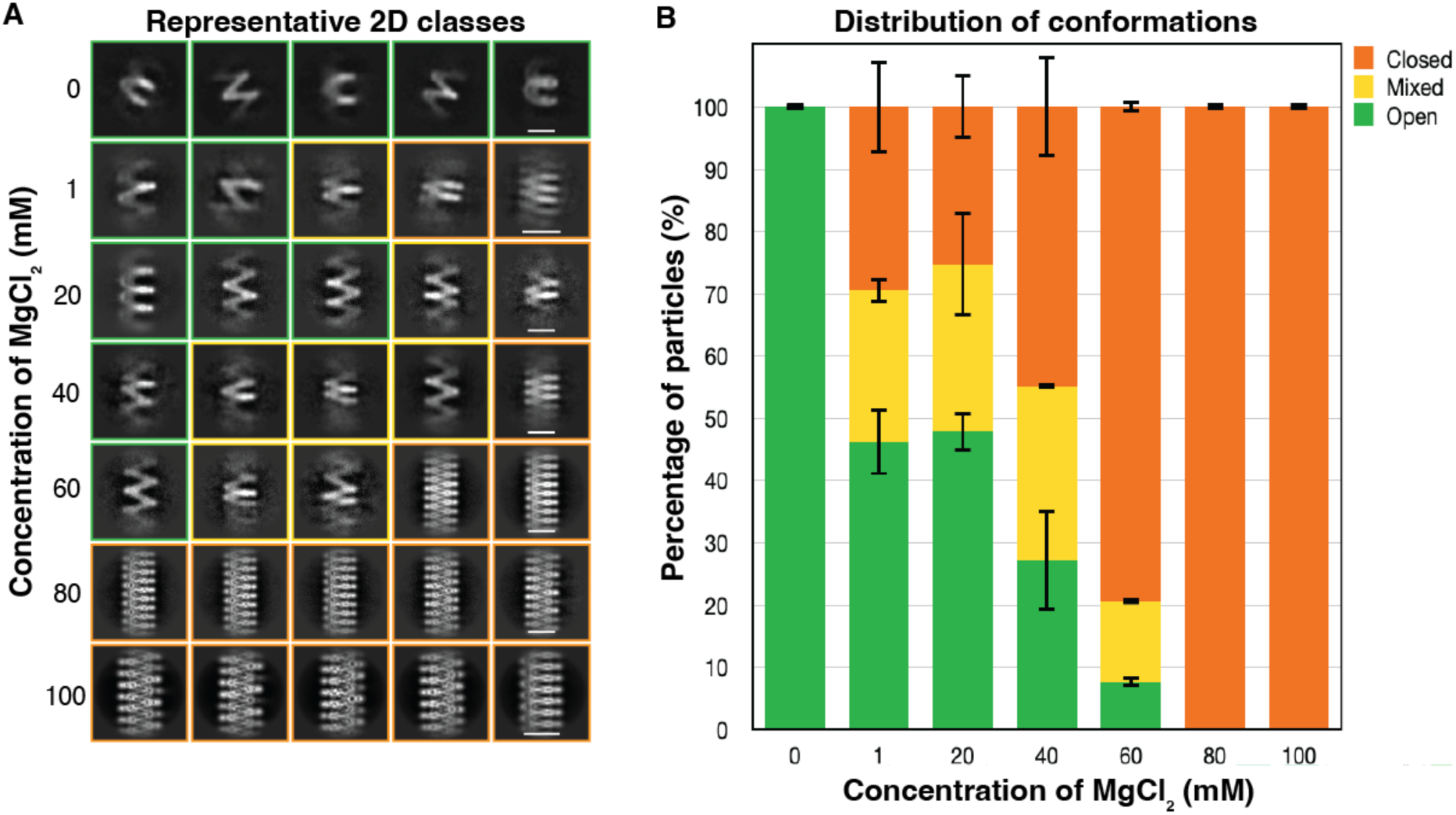
Effect of MgCl₂ concentration on the distribution of HHoB-DNA complex conformations. (A) Representative 2D class averages from particle picking using the same Topaz models across all datasets. Enclosing boxes indicate open (green), mixed (yellow), and closed (orange) conformations. Scale bar = 10 nm. (B) Stacked column plot showing the percentage distribution of conformations in each dataset. Error bars represent the standard deviation from technical duplicates, obtained using two independent Topaz models for particle picking.

Interestingly, even at low Mg²⁺ concentrations (1 mM and 20 mM), individual nucleosomes stacked with each other to form extended hypernucleosomes (Figure 2A). These hypernucleosomes appeared in various configurations: completely open, closed, or "mixed". With increasing Mg²⁺ concentration, hypernucleosomes of increasing length were formed (Figure S4). The hypernucleosomes exhibited very regular arrangement and spacing, and were consistently straight. The spacing (superhelical pitch) in the closed hypernucleosomes was 24.6 Å, while in the open configuration it was 63 Å, consistent with the spacing of individual nucleosomes in the 1 mM Mg²⁺ condition. The pitch of the closed hypernucleosome was smaller (24.6 Å) than in the individual closed nucleosomes (29.5 Å), where the DNA ends are more flexible, as they are not packed into a rigid hyper-assembly. EMSA performed at 10 mM Mg²⁺ concentrations confirmed that the cooperativity of HHoB-binding to DNA increased, and instead of the ladder-like shifts in EMSA, HHoB binding resulted in a single shifted band similar to HMfB (Figure S1). Histones HHoF and HHoG exhibited the same behavior (Figure S1).

Hypernucleosome formation is likely mediated by Mg²⁺ ions binding to the negatively charged phosphate DNA backbone in a non-site-specific manner, as we could reproduce the effect of hypernucleosome formation in the “high Mg” (100 mM) condition by substituting Mg²⁺ with other divalent cations – Zn²⁺ and Ca²⁺ (Figure S5), and as we could not observe any defined coordinated binding sites near the protein density (although the high resolution would allow us to resolve such sites). We note that Mg²⁺ is the most abundant divalent ion in the cells^34^, with concentrations typically 100-1000 times higher than Ca²⁺ or Zn²⁺, thus influence of other divalent cations compared to Mg²⁺ would be, in principle, negligible in native conditions.

We then tested complete absence of Mg²⁺ ions during nucleosome reconstitution – and acquired a cryo-EM dataset in the "zero Mg" condition. Single particle analysis revealed only one population of particles – all nucleosomes were in the open conformation (Figure S6, Video S1). In this condition all nucleosomes existed as individual particles, and formation of hypernucleosomes was not observed.

### Cryo-EM Analysis Reveals the Structures of the Closed and Open HHoB Hypernucleosomes

To determine the molecular structure of the closed hypernucleosome at high resolution, we collected a large cryo-EM dataset on the 100 mM Mg²⁺ sample, where 100% of particles were in the closed conformation. Closed hypernucleosomes formed true helical structures, which were processed accordingly in cryoSPARC (Figure S7). We resolved the map at 2.6 Å reported resolution (Figure 3B, Figure S7, Table 1, Video S1). The high resolution of the map allowed us to place side-chains with high confidence during model building. The closed hypernucleosome structure (PDB 9QV6) is similar to the HMfB hypernucleosome^6^ (RMSD of 0.82 Å over 3 histone dimers at Cα atoms). Closed conformations from the 1 mM and 100 mM conditions were very similar (RMSD 0.87 Å over 3 histone dimers at Cα atoms). HHoB dimers are continuously bound to the DNA, wrapping it into a tight left-handed super-helix. Individual DNA molecules (147 bp) within the assembly are "seamlessly" connected via DNA end-to-end contacts, and the histone dimers are continuously bound to the DNA with a footprint of ∼30 bp.

**Figure 3.**
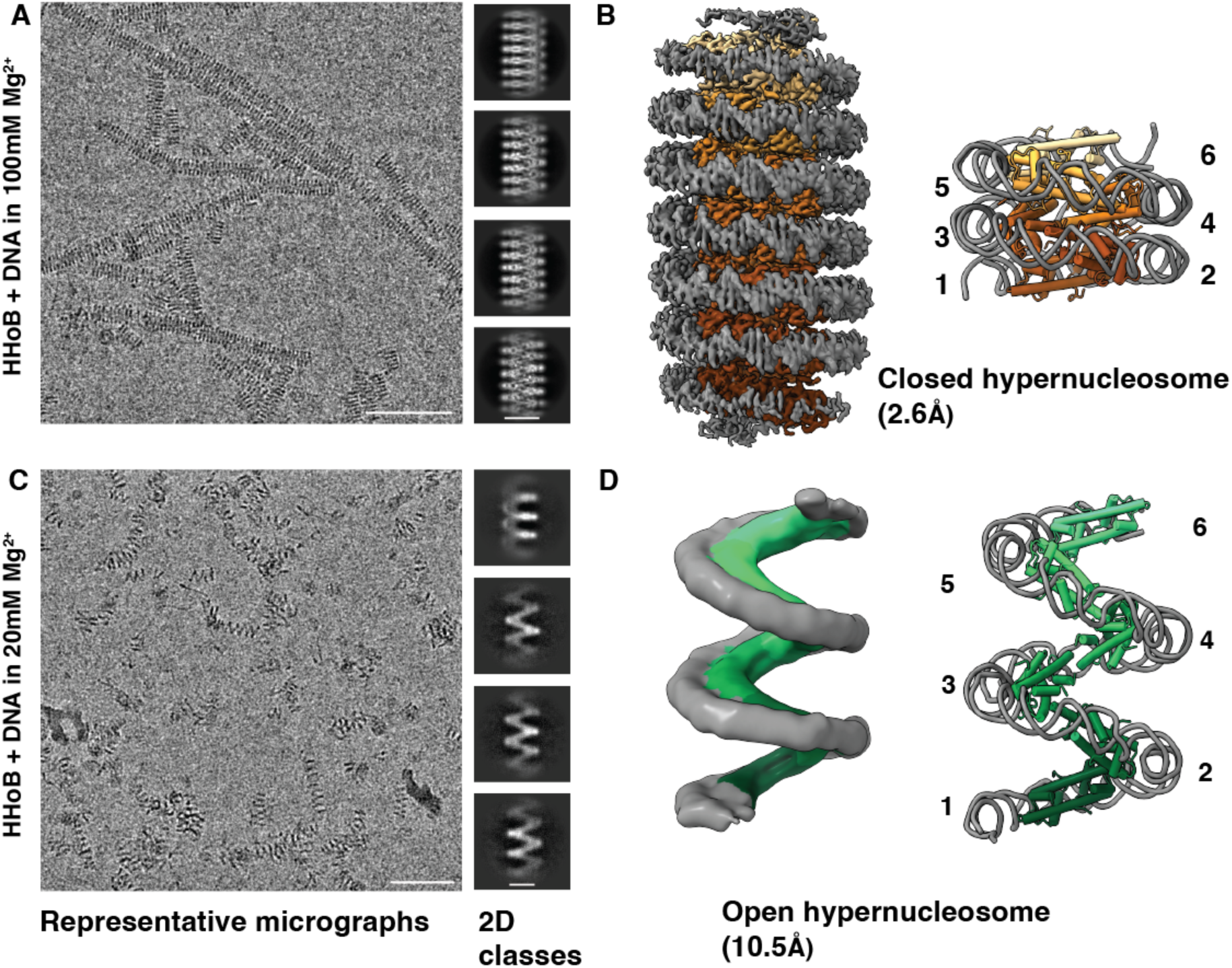
HHoB hypernucleosomes in open and closed states. (A) Representative micrograph (left, scale bar = 50 nm) and 2D class averages (right, scale bar = 10 nm) of the HHoB–Widom601 147 bp complex in 100 mM MgCl₂ buffer. (B) Refined EM map (left) and molecular model (right) of the HHoB closed hypernucleosome (PDB 9QV6), with 180 bp of DNA in grey and six histone dimers in shades of orange. (C) Representative micrograph (left, scale bar = 50nm) and 2D class averages (right, scale bar = 10nm) of the HHoB–Widom601 147 bp complex in 20 mM MgCl₂ buffer. (D) EM map (left) and rigid-body fit of two HHoB open conformation models (right), with 180 bp of DNA in grey and six histone dimers in shades of green. Scale bar 50 nm.

Next, we collected a cryo-EM dataset on the 20 mM Mg²⁺ condition sample to enrich for the open hypernucleosome state. The open hypernucleosomes exhibited greater flexibility compared to the closed state, and consequently were resolved at a lower resolution of ∼10 Å (Figure 3C-D, Figure S8, Video S1). The open hypernucleosome conformation is similar to the individual open nucleosome conformation from the 1 mM condition. We fitted the open nucleosome model built from our 3.6 Å map (1 mM condition) as a rigid body into the open hypernucleosome map with high confidence (Figure 3D). In both open and closed hypernucleosomes, the DNA ends interact to form a continuous superhelix, which is wrapped along its length by histone dimers (Figure 3C-D). In the open hypernucleosomes, the “side- way” contacts between histone dimers and the DNA end-to-end stacking are enough to drive the assembly formation, even in the absence of histone stacking interactions as in the closed form.

To exclude a possible influence of the SELEX-based Widom601 DNA sequence on hypernucleosome formation, we next used a native genomic 420 bp DNA sequence (derived from a HeimC3_31310 gene of LC3 metagenome) for reconstitutions. The EMSA showed typical binding behavior comparable to the Widom601 DNA (Figure S2). The number of shifted bands was higher than for the 147 bp Widom DNA due to the greater number of binding sites on the longer native DNA. Importantly, in the “high Mg” (100 mM) condition, by cryo-EM we observed formation of hypernucleosomes similar to those formed with Widom601 DNA (Figure S5). The pitch of the native-sequence hypernucleosome was measured to be 25.2 Å – matching that of the closed hypernucleosome formed on Widom601 DNA (25.4 Å).

### The Closed and Open Nucleosomes Differ in Key Interfaces

Three key interfaces are important for nucleosome and hypernucleosome assemblies: 1) histone dimer-DNA interface; 2) histone dimer-dimer interface; and 3) histone dimer stacking interface. Another important contact is the histone-histone interface within the dimer "handshake" motif, but this is extremely conserved across the phylogenetic tree^4^ and is essentially identical between the assemblies (RMSD 0.47 Å over the dimer between Cα atoms).

The open state of the HHoB hypernucleosome forms a highly extended superhelix without any stacking interactions between histones (Figure 4). This extremely open conformation is fully sustained by the histone-DNA interfaces and minimal histone dimer-dimer interfaces (Figure 4). Each histone dimer in the open state participates in two dimer-dimer contacts (558 + 526 Å² interfaces) and one histone-DNA interface (1966 Å^2^ area). In the closed hypernucleosome state, each histone dimer is involved in two dimer-dimer interfaces (727 + 728 Å^²^), one histone-DNA interface (1959 Å^2^), and two stacking interfaces (1674 + 1676 Å^2^). Overall, the interface area that a histone dimer participates in within the closed hypernucleosome is 2.2x larger (6764/3050 = 2.2) than in the open conformation, making it much more compact and stable.

**Figure 4.**
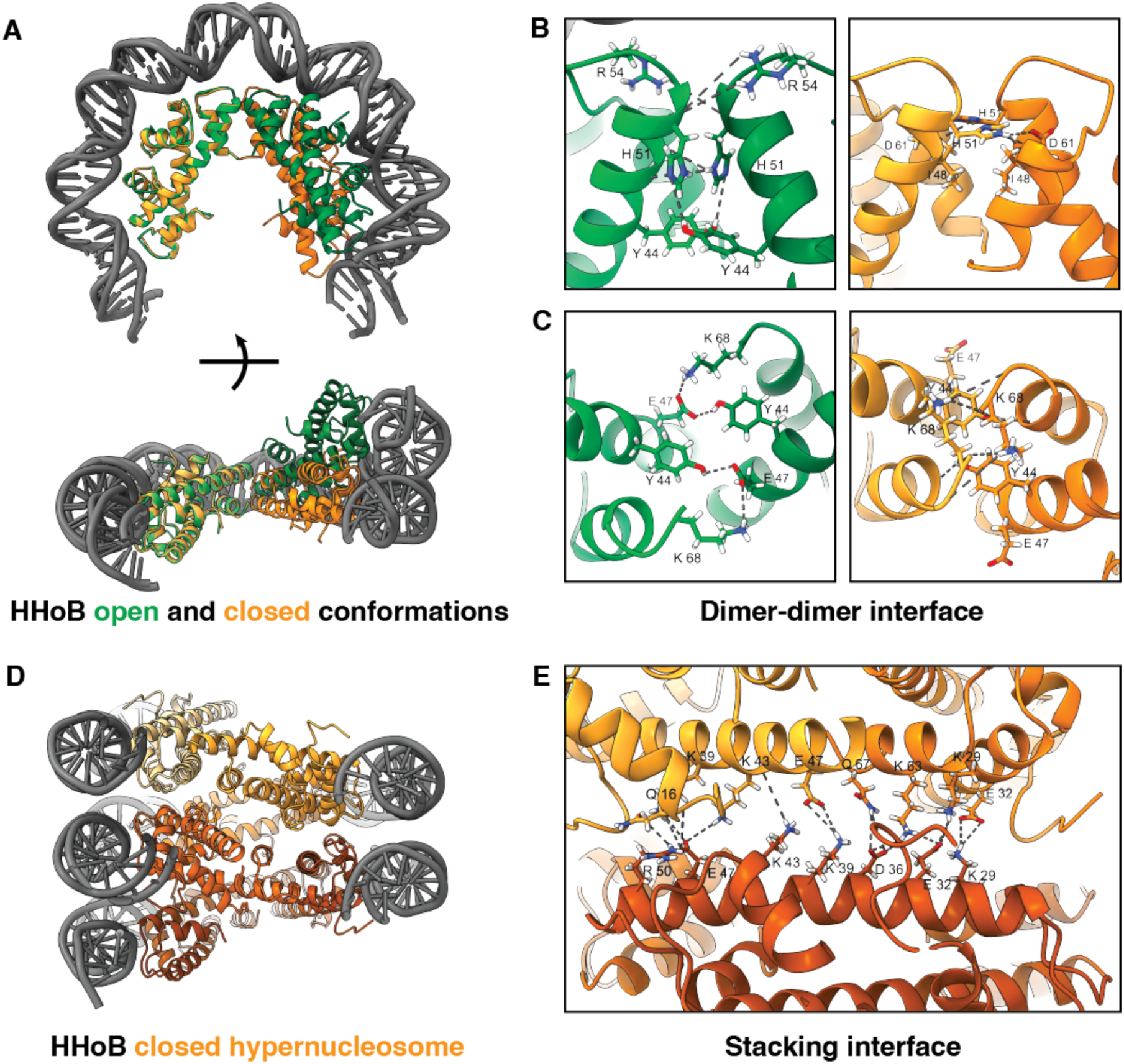
Key interfaces involved in the open (shades of green) and closed (shades of orange) conformations of the HHoB - DNA complex. DNA is colored in grey. (A) N and N+1 histone dimers of open (PDB 9QV7) and closed (PDB 9QV5) conformations, aligned along one histone dimer. (B) and (C) show key residues and interactions involved at the dimer-dimer interface: Y44, E47, I48, H51, R54, D61, K68. (D) Slice view of the HHoB closed hypernucleosome (PDB 9QV6) and (E) shows the stacking interface between N, N+2 and N+3 histone dimers. Key residues involved are Q16, K29, E32, K39, K43, E47, R50, K63, Q67. Dashed lines indicate electrostatic and hydrogen bond interactions.

The histone-DNA interface is highly conserved and is essentially identical in the open and closed nucleosome states reported here (RMSD 0.40 Å between Cα atoms) (Figure 4). The main residues involved are Arg9, Arg15, Arg21, Lys55, Lys58 (Figure 1A).

### Histone Dimer-Dimer Interface Changes Drastically Between the Open and Closed States

Comparison of the histone dimer-dimer interface in the open and closed nucleosome models revealed substantial rearrangement between states, with an RMSD between Cα atoms of a dimer N+1 of 12.95 Å (if dimers N of the open and closed state are superimposed) (Figure 4A, Video S2). The relative positions of the two neighboring dimers differ strongly in the closed vs open state. In the closed state the dimers are positioned at a shallow angle to one another (Figure 4A, model in orange), allowing for a gentle rise of the helix in a closed hypernucleosome. While in the open state the neighboring dimers are positioned at a steeper angle to one another (Figure 4A, model in green), therefore the rise of the superhelix is also much steeper allowing for a very open extended structure. The dimer in the open structure is by 21.8 degrees at a more steep angle to the first dimer compared to the closed state (measured between α2 HHoB helices in ChimeraX^35^).

Residues Tyr44, Glu47, Ile48, His51, Arg54, Asp61, and Lys68 are involved in interactions between helices α2 and α3 and loop L2 of the interacting histones in the open state (Figure 4B- C). In the closed state, residues Leu64, and Gln67 additionally contribute to the dimer-dimer interface, and many of the above-mentioned residues change their interaction network. In the open state, the interface area of dimer N with dimer N-1 is 558 Å² compared to 727 Å² in the closed state. Moreover, the His51 and Tyr44 side chains are oriented very differently in the closed and open states (Figure 4B-C). Residue His51 is highly conserved (with a few exceptions), while others are less conserved. Interactions Tyr44-Glu47, Tyr44-His51, and Glu47-Lys68 are unique to the open state.

Interestingly, the eukaryotic “octasome” formed by four H3-H4 heterodimers also shows an open clam-shell-like conformation^36^. The opening (equivalent to ”pitch”) of the octasome open state is 53 Å – much closer to the HHoB open state (63 Å) than to the closed one (25 Å). Notably, at the central H4-H4 histone interface in the octasome structure, there is also a Tyr residue but at an equivalent of position 48 (HHoB numbering). We hypothesized that if the Tyr helps to stabilize the open state of the HHoB dimer-dimer interface, its exact position on the α2 helix may define the final degree of nucleosome opening. Therefore, we generated a double mutant (“mut1“ Y44A-I48Y) where the original Tyr44 was mutated to Ala, and instead a Tyr was introduced at a position 48 to imitate the Tyr position in the H3-H4 octasome. The mutant was expressed and purified using the same protocol as the wild-type HHoB. It showed similar DNA binding behavior as the wild-type HHoB via EMSA (Figure S2). This was expected, as the histone-DNA interface was not affected in the mut1. We then collected a cryo-EM dataset on the nucleosomes reconstituted from mut1 HHoB in the 1mM Mg condition, and analyzed the data via SPA (Figure S9). In the 2D classes we observed both closed and open states (Figure 5F), although interestingly, the nucleosome opening in the open classes was smaller than in the wild-type HHoB. The 3D reconstruction resulted in a low-resolution map, indicating that the destabilization of the dimer-dimer interface weakens the entire nucleosome conformation, or leads to higher heterogeneity of nucleosome conformations. The final average EM map shows a decreased nucleosome opening compared to the wild-type HHoB open nucleosome (Figure 5F, S9). The degree of nucleosome opening in mut1 open state matched most closely the H3- H4 octasome structure, which has a Tyr in a matching position 48 (Figure S9). This supports our hypothesis regarding the role of Tyr44 in open state stabilization. Notably, all residues involved in the open HHoB nucleosome dimer-dimer interface are also involved in the closed dimer-dimer interface formation, therefore they cannot be completely decoupled in a mutagenesis experiment.

**Figure 5.**
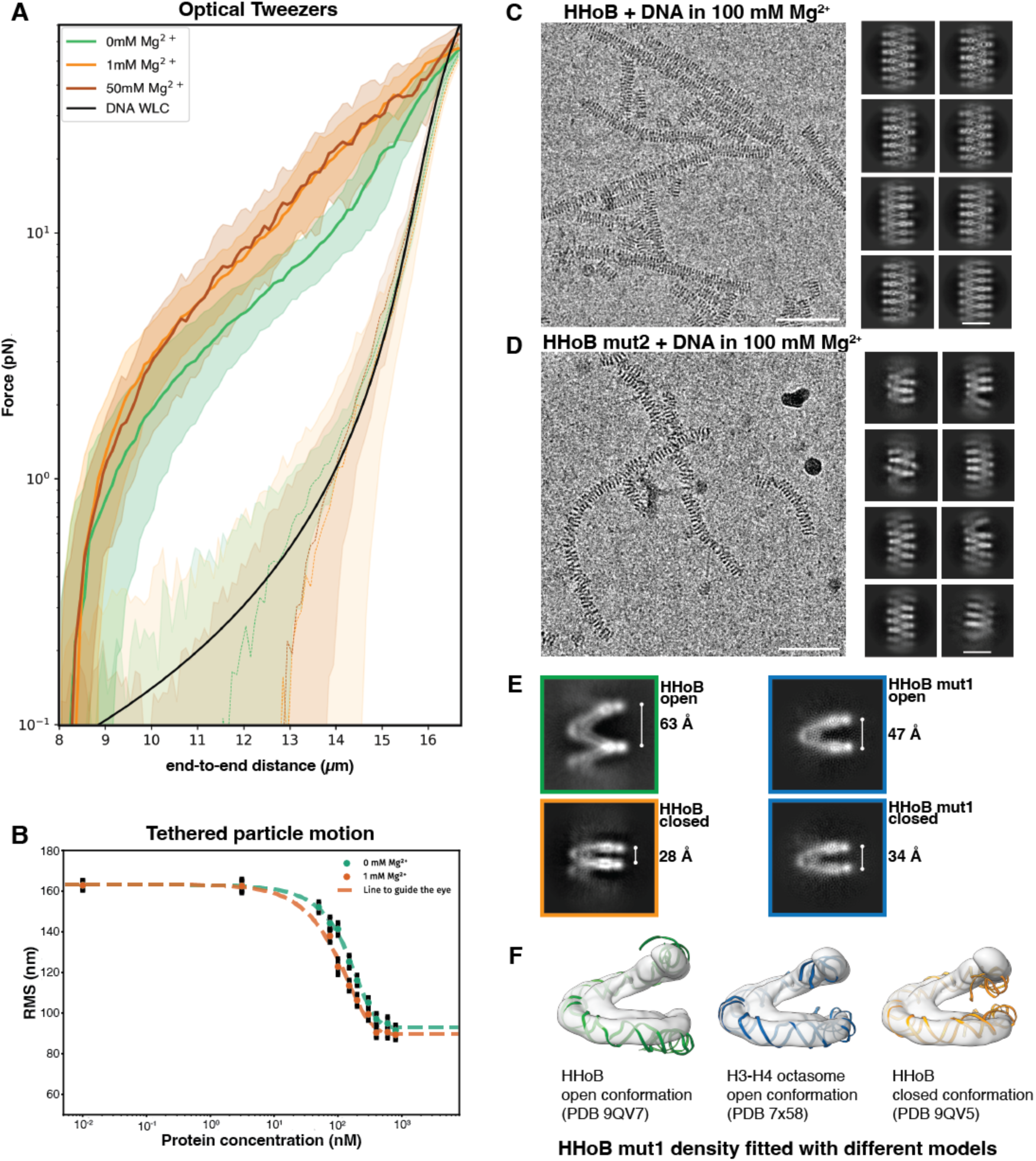
Biophysical characterization of the HHoB–DNA complex in the presence of magnesium ions. (A) Force- extension curves of HHoB–Lambda phage DNA (48.5 kb) in 0mM, 1 mM and 50 mM MgCl₂ (green, orange, and brown, respectively). Solid lines and shading indicate the mean and standard deviation (n=10). The black solid line represents the theoretical Worm-Like Chain (WLC) for DNA at 25°C. (B) TPM experiments of HHoB with 685 bp DNA at varying Mg²⁺ concentrations. Measurements were performed in duplicates; error bars indicate standard deviation. (C, D) Representative micrographs (scale bar 50nm) and 2D class averages (scale bar 10nm) of the HHoB–DNA complex and HHoB mut2–DNA complex in 100 mM MgCl₂. (E) 2D classes of HHoB mut1 – DNA in 1 mM MgCl₂ (blue) are shown alongside corresponding view from open (green) and closed (orange) conformations of the wild type complex. The superhelical pitch was intermediate. (F) A low-resolution density map of the HHoB mut1–DNA complex (transparent grey) was fitted with DNA models from the open (green) and closed (orange) wild type molecular models, as well as the open conformation of the H3-H4 octasome (blue).

### Stacking Interface in the Closed HHoB Hypernucleosome is Mediated by Electrostatic Interactions

Another important contact that stabilizes the closed state is the histone "stacking" interface. Based on the closed hypernucleosome structure, stacking interactions are primarily mediated by electrostatic interactions between α3 and α2 helices of histone dimer N1 and the α2 helix of dimer N3, and between N1 α2 and α1, L1 of dimer N3. The key residues involved in stacking interactions based on our structure are: Gln16, Lys29, Glu32, Asp36, Arg50, and Lys63 (Figure 4E). The exact positions of these residues are not completely conserved between HMfB and most short LC3 histones (Figure 1A), but in both structures they create a "velcro-like" arrangement of positively and negatively charged residues that mediates the stacking interface.

We generated a triple mutant “of the key residues (“mut2” K29A E32A D36A) involved in the stacking interface and tested the ability of the mutant to form nucleosomes and hypernucleosomes. EMSA results showed that the triple mutant has similar DNA binding properties to wild-type (Figure S2), as expected since the histone-DNA interface remains unaffected by these mutations. We then tested the ability to form hypernucleosomes in a high Mg²⁺ condition (100 mM) and examined the samples by cryo-EM (Figure 5E). We observed that the stacking interface mutant can form hypernucleosomes (as not all stacking interactions are disrupted in a mutant), but they are much less ordered than wild-type, with a more flexible and "wobbly" appearance (Figure 5D-E, Figure S10). The SPA analysis enriches for more ordered states, and therefore does not reflect the whole range of conformations in the mutant nucleosomes. Nonetheless it reveals a much larger pitch compared to the wild-type closed hypernucleosomes (Figure S10). This demonstrates that the stacking interface is essential for stabilizing the closed hypernucleosome state.

### Biophysical Experiments Show Distinct Behavior of HHoB Nucleosomes Depending on Mg²⁺ Concentration

To investigate the mechanical properties of the HHoB hypernucleosome in response to Mg²⁺, we performed optical-tweezer-based force-spectroscopy measurements^37,38^ using long biotinylated Lambda phage DNA (48,502 bp) as a substrate. The Mg²⁺ concentration served as a proxy for chromatin compaction, with cryo-EM data indicating that, in the absence of Mg²⁺, the histone-DNA complex exists exclusively in an open conformation. In contrast, samples containing Mg²⁺ exhibit a mixture of open and closed conformations. Starting at 80 mM Mg²⁺, only the closed conformation is observed. Force-extension measurements of the HHoB-DNA complex were performed in buffers containing 0 mM, 1 mM, and 50 mM MgCl₂ (Figure 5A). These curves represent the mean ± standard deviation from 10 independent force-extension traces. The concentration of HHoB mut3 (a fluorescently tagged mutant of HB, see methods) in the protein channel was 200nM, which is 6.5 times higher than the measured Kd (30.8 ± 3.7 nM). We overlay the theoretical Worm-like Chain (WLC) model for free DNA (Figure 5A) which correlates very well with the free DNA curves from the experiments. The free DNA curves also demonstrate no difference in mechanical properties of DNA in the presence of magnesium ions. Notably, the overall force-extension curves resemble previously published data on histone HMfB hypernucleosomes^9^. In the presence of HHoB histone, the force magnitudes measured for the 0 mM Mg²⁺ condition are distinct from those obtained at 1 mM and 50 mM MgCl_2_ along the extension range (Figure 5A). To quantify these differences, we probed the force difference between histone-DNA complexes and free DNA at low (10 μm) and high (13 μm) end-to-end distances (EED). At low EED, the measured force differences were 2.01 ± 0.36 pN for 0 mM, 3.52 ± 0.59 pN for 1 mM, and 3.27 ± 0.68 pN for 50 mM MgCl₂ (N = 10). While at high EED, the force differences were 6.63 ± 0.56 pN for 0 mM, 12.73 ± 2.03 pN for 1 mM, and 13.41 ± 1.94 pN for 50 mM MgCl₂ (N = 10). Paired t-tests confirmed that the differences between 0 mM and both 1 mM and 50 mM MgCl₂ were statistically significant at both EEDs (p < 0.01). It is worth noting that due to clogging issues in the microfluidic flow channels at higher protein concentrations, our experiments are limited in probing the full saturation regime of histone binding to DNA. However, clearly, the consistent and significant force differences across conditions suggest a genuine Mg²⁺- dependent modulation of DNA compaction by HHoB, possibly arising from hypernucleosome stacking interactions, cooperative histone binding, or a combination of both.

To further investigate the structural changes induced by HHoB upon binding to DNA in solution, we conducted Tethered Particle Motion (TPM) experiments using a linear 685 bp dsDNA fragment. Importantly, TPM experiments allow us to probe full range of protein concentrations up to the full saturation regime. In TPM, a bead is tethered to a glass surface by a DNA molecule^39^. DNA compaction can be quantified by measuring the decrease in root mean squared (RMS) displacement of the bead relative to the glass surface^9,40^. Individual data points represent the average of N>100 protein-DNA complexes. Titration of HHoB induced a gradual decrease in RMS, confirming that HHoB wraps and compacts the DNA (Figure 5C). Next, we examined the effect of Mg²⁺ in an otherwise identical protein titration experiment. These experiments showed that in the presence of Mg²⁺, hypernucleosome formation shifts toward lower protein concentrations, and the RMS of the hypernucleosome structure obtained at protein saturation was more compact than in its absence (Figure 5C). We note that when the DNA template is longer (685 bp in TPM experiments), the hypernucleosomes will also be able to form at lower Mg concentrations, while stacking of individual nucleosomes needs higher Mg concentrations (in our cryo-EM experiments). These biophysical experiments provide direct evidence that Mg²⁺ modulates hypernucleosome mechanical properties to form a more compacted state, as higher forces are needed to disrupt the chromatin fiber in the presence of Mg²⁺ than in its absence, and is in perfect agreement with our cryo-EM observations.

## Discussion

### Closed Nucleosome State is Conserved Across Archaea

This study provides the first structural insights into Asgard chromatin, expanding our understanding beyond the better-studied Euryarchaeota^6,10,41,42^. Asgard archaea, and Hodarchaea in particular^13^, are the closest living archaeal relatives to Eukaryotes known to date, which makes studying their genome organization extremely important in the context of evolution. Here, we characterized the tail-less histone HHoB from LC3 Asgard Hodarchaea and demonstrated that it forms nucleosomes in two distinct conformations – closed and open.

All previously reported archaeal nucleosome structures were in a closed conformation (Figure S11)^6,10^. This includes the classical closed nucleosomes formed by Euryarchaeal HMfB^6^ and HTkA^10^, as well as the “slinky” arrangement of HTkA^10^, where histones pack into two closed nucleosomes which are connected at an angle to each other by a short linker DNA. Histone dimer-dimer and stacking interfaces in the slinky^10^ and classical^6^ Euryarchaeal nucleosome structures are consistent with the HHoB closed state. Since both HMfB/HTkA and HHoB histones can form closed nucleosomes, this closed conformation appears to be conserved across distant archaeal groups, such as Asgard and Euryarchaeota. Our findings also experimentally validate the earlier prediction^8^ that HHoB LC3 can form closed hypernucleosomes, engaging interactions at the stacking interface. Notably, the open Asgard HHoB nucleosome conformation is novel and was not predicted in previous studies.

### Open Hypernucleosomes as an Asgard Innovation

The HHoB open state reported here represents a distinct structural configuration of archaeal chromatin. The HHoB open state has a different dimer-dimer interface compared to the closed state (and to all structures reported previously^6,10^) and is characterized by a complete absence of histone stacking interfaces.

Our open HHoB nucleosome structure is characterized by key interactions between Tyr44- Glu47 and Tyr44-His51. All three amino acids are conserved in six out of ten LC3 histones, except in HHoA, HHoC, HHoD, and HHoF. Overall, the key residue Tyr44 is present in 11% of known Asgard histones^11^ and in none of the annotated Euryarchaeal histones (examples in Figure S12). This suggests that the open nucleosome state could be a unique innovation of Asgard archaea and may be relatively widespread among them. Importantly, a eukaryotic nucleosome assembly “H3-H4” octasome displays an “open” state as well^36^. It was suggested that the H3-H4 assembly might be ancestral^43^ to the canonical nucleosomes. In that light, it is striking, that the Asgard open nucleosome resembles the conformation of the potentially ancestral eukaryotic H3-H4 assembly, indicating the open conformation as an intermediate step in nucleosome evolution (see comparison in Figure S11). However, more experimental evidence of open nucleosome examples is needed to more clearly define the sequence features responsible for this conformation. Notably, AlphaFold3^44^ is unable to predict structures in the open conformation even for the HHoB histone, potentially due to training bias, but instead predicts a closed conformation for all.

We speculate that the open nucleosome state may confer particular advantages for Asgard archaea, especially in enabling faster and more efficient histone variant exchange compared to the highly compact closed assembly. Recent analyses have shown that Asgard genomes often encode a larger number of canonical “nucleosomal”^45^ histone variants than many Euryarchaea, which typically encode only one or two (e.g., HMfB/A, HTkA/B)^11,46^. To exchange a histone dimer even at the end of a closed hypernucleosome, numerous interfaces must be disrupted (stacking, dimer-dimer, and dimer-DNA), whereas in the open assembly, the stacking interface is already absent, thus making histone dimer exchange less energetically costly. Additionally, the open chromatin state may minimize steric clashes involving histones with extended structural elements, such as additional N- or C-terminal α-helices or tails – features more frequently observed in Asgard histones compared to Euryarchaea – thus facilitating the “evolutionary exploration” of histone extensions in Asgard archaea.

The presence of Asgard open hypernucleosomes also suggests a stable yet dynamic chromatin structure, where DNA is bound by histones but both remain accessible for chromatin modification factors – potential chromatin “readers” and “writers” predicted in Asgard archaea^47^. The open state also could be easier for the DNA-machinery to passage through than in the closed state, based on the lower number of contacts that need to be broken. In contrast, closed hypernucleosomes, may partially limit the accessibility of histones and DNA more than the open hypernucleosomes, suggesting a regulatory interplay of the two states. This balance of stability and accessibility could be particularly beneficial for thermophilic and hyperthermophilic Asgard species, which must stabilize their DNA under extreme conditions via histone association while maintaining accessibility^48^. Interestingly, the last common ancestor of Asgard archaea is suggested to have been a thermophile^49^.

### Mechanisms of Mg²⁺-Based Nucleosome-State Regulation

We investigated the effect of Mg²⁺ ion concentrations on chromatin structure, as Mg²⁺ plays a crucial role in nucleosome state regulation in both archaea and eukaryotes. We found that, within a wide range of Mg²⁺ concentrations (1–60 mM), Asgard histone HHoB forms both closed and open assemblies. At higher Mg²⁺ concentrations, HHoB leads to the formation of long closed hypernucleosomes.

Our results suggest a possible mechanism for Mg²⁺-dependent regulation of archaeal chromatin structure. Many archaea use a "salt-in" mechanism^50^ to adjust intracellular ion concentrations based on extracellular levels. In this context, Mg²⁺ ions could regulate chromatin state by binding to the negatively charged DNA phosphate backbone, shielding charges, and stabilizing DNA folding. At higher Mg²⁺ concentrations, DNA gyres in hypernucleosomes may move closer together, promoting a closed state that stabilizes protein-protein interactions at both the histone dimer-dimer and stacking interfaces. We further demonstrated through EMSA assays that the cooperativity of HHoB histone-DNA binding is generally much lower compared to Euryarchaeal HMfB in zero Mg²⁺ conditions. Notably, the cooperativity of HHoB binding increases with higher Mg²⁺ concentrations, correlating with increased formation of closed hypernucleosomes. This suggests that Mg²⁺ may play a role in regulating chromatin states through its effect on histone-DNA binding cooperativity. Our results suggest that the cooperativity of HHoB binding to the DNA is defined by the synergy of three factors: Mg shielding DNA charges, formation of dimer-dimer and stacking interfaces.

### Asgard Chromatin Model

The current model of archaeal chromatin may be described as "variable beads on a string," where in each nucleosome, a variable number (N) of histone dimers associates with DNA, wrapping it into hypernucleosomes of 30×N length. Our work suggests that this model can now also be applied to describe Asgard chromatin. We observe the formation of long stable hypernucleosomes in both closed and open states, formed by just one histone variant out of ten encoded in the genome. Taking into account the presence of nine other types of histones in the LC3 metagenome, we expect both closed and open hypernucleosomes of variable length to be formed locally by one or several types of histones (model in Figure 6). Meanwhile, other types of histones (with lower propensity to form hypernucleosomes) as well as nucleoid-associated proteins (NAPs) could act as capstones^51^ and road-blocks, respectively, thus regulating the size and spread of hypernucleosomes and, in turn, DNA accessibility. The prevalence of open versus closed conformations could be regulated either by Mg²⁺ concentration, or by other local environment factors (cations, polycations, highly charged small molecules), or stabilized by additional protein players. Overall, further extensive structural and functional studies are required to understand the full conformational landscape of Asgard chromatin, in order to shed light on the role of multiple histone variants, including histones with tails, and how they can shape and regulate chromatin organization.

**Figure 6.**
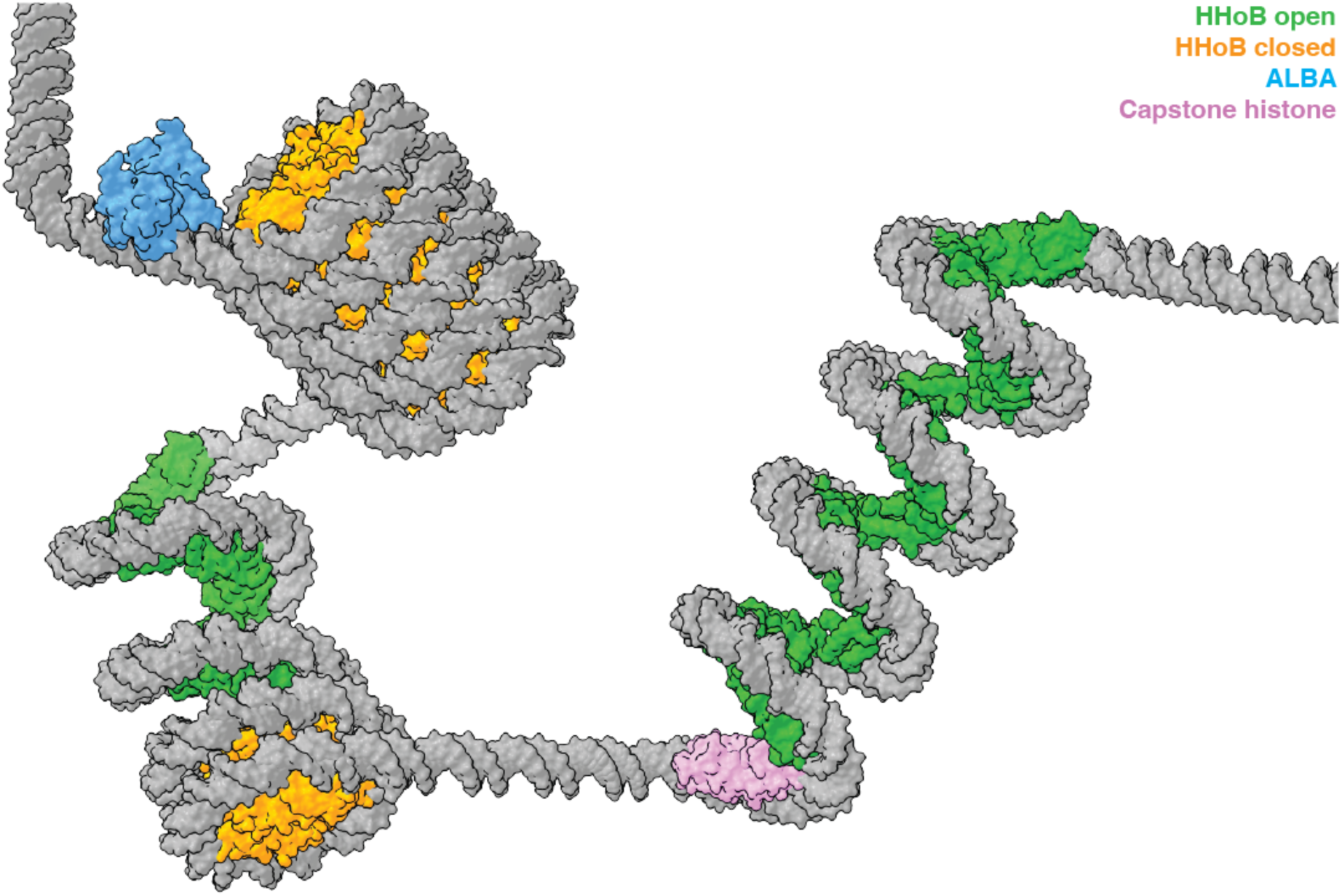
Asgard ‘variable beads-on-a-string” chromatin model. Hypernucleosomes of varying lengths either in open (green) or closed (yellow) conformations. Other histone variants (pink) and nucleoid associated proteins like Alba (blue) could act as capstones and roadblocks.

### Resource availability

#### Lead contact

Correspondence and requests for materials should be addressed to SOD.

#### Materials availability

Materials are available from Svetlana Dodonova upon request under a material transfer agreement.

#### Data and code availability

The cryo-EM density reconstructions and model coordinates were deposited with the Electron Microscopy Database (EMDB) and Protein Data Bank (PDB) under accession codes: EMD- 53386 and PDB 9QV5 (closed nucleosome in 1 mM Mg^2+^), EMD-53388 and PDB 9QV7 (open nucleosome in 1 mM Mg^2+^), EMD-53390 (open nucleosome – 0 Mg^2+^), EMD-53387 and PDB 9QV6 (closed hypernucleosome), EMD-53389 (open hypernucleosome).

This paper does not report original code.

Any additional information required to reanalyze the data reported in this paper is available from the lead contact upon request.

## Acknowledgments

This work is supported by the European Union (ERC Starting Grant 3DchromArchaea, grant agreement no. 101076671, to SOD). HMR acknowledges support by the EMBL International PhD program. RTD was supported by the Dutch Research Council OCENW.GROOT.2019.01. We thank Wim Hagen, Felix Weiss, Joseph Bartho and Sebastian Unger (EMBL MSB cryo-EM specialists) for technical assistance. We thank Alexander Marchanka for help with mutagenesis and initial optimization. We thank the EMBL PEP core facility and Karine Lapouge for support with protein quality control experiments. We thank Christoph Müller, and Fredrika Rajer for helpful comments on the manuscript.

Funded by the European Union. Views and opinions expressed are however those of the authors only and do not necessarily reflect those of the European Union or the European Research Council Executive Agency. Neither the European Union nor the granting authority can be held responsible for them.

## Author contributions

HMR prepared the DNA and protein constructs, purified the proteins and DNA, carried out in vitro reconstitutions, EMSA experiments, prepared cryo-EM grids, collected and processed cryo-EM data, built the structural models. HMR together with NLB performed the force-extension experiments. MKMC designed and carried out the footprinting EMSA and TPM experiments under supervision of RTD. HMR with help from TQ analyzed force-extension data. SOD conceived the study, supervised the research and wrote the manuscript, with input from all authors.

## Supplemental information

Document S1 - Materials and Methods

Supplementary Figures S1-S12

Video S1. Gallery of EM maps and corresponding models, related to Figure 1 and 3.

Video S2. Conformational change between the histone dimer-dimer interface in the closed and open states. Related to Figure 4.

## Supplementary Materials

### Materials and Methods

#### Plasmids and strains

Full-length histone sequences from Asgard genome Hodarchaeaota LC_3 (GCA_001940645.1) were codon-optimised for expression in *E. coli*, and then incorporated into the LIC 1B plasmid from MacroLabs (pET His6 TEV LIC cloning vector 1B - Addgene #29653). Accordingly, the HHoB histone (Uniprot A0A1Q9NRY6) expression construct in a LIC 1B vector contained an N-terminal 6×His-tag followed by a tobacco etch virus (TEV) protease cleavage site.

Histones HHoF (Uniprot A0A1Q9N8N6), HHoG (Uniprot A0A1Q9NAM9) and HMfB from *Methanothermus fervidus* (Uniprot P19267) were also incorporated into LIC 1B expression plasmids.

#### HHoB mutagenesis

Mutants of HHoB were generated using PCR-based mutagenesis. The specific mutations introduced were as follows: HHoB mut1(Y44A I48Y); HHoB mut2 (K29A E32A D36A); and HHoB mut3, which included a wild-type HHoB with additional GGGC at the C-terminus (later used for labelling). Constructs for mut1 and mut2 were ordered from IDT as part of a pET-IDT expression vector. For mut3, following PCR amplification, the mutant plasmids were transformed into *E. coli* XL-blue cells and screened by colony PCR. Plasmid DNA was extracted using a Miniprep Kit (Qiagen) and the mutations were validated by Sanger sequencing (Eurofins Genomics).

#### Protein expression and purification

HHoB, mut1, mut2 and mut3 proteins were expressed in *E. coli* BL21-CodonPLus (DE3)-RIL strain by induction (0.5 mM IPTG at OD_600_ 0.5-0.6) and incubated at 37°C for 3-4 hours. Cell pellet was harvested by centrifugation at 4000 rpm at 4°C in and stored at -20℃. Cell pellets were thawed in lysis buffer containing 20 mM HEPES pH 7.5, 300 mM NaCl, 5% glycerol, 1% Triton X-100 and 1x protease inhibitor (PI tablet, Roche) for 20 min at 4°C. Lysis was further carried out by sonication for 10 min at 30% power and 0.4 s duty cycle using a Branson sonifier. The cell lysate was centrifuged at 27,000 g for 20 min at 4℃. The supernatant was collected, and His-tagged histone HHoB was enriched using affinity purification using ROTI®Garose-His/Ni NTA-HP beads (Carl Roth). The His-tag was cleaved off using GST-tagged TEV protease during overnight dialysis into buffer B (20 mM HEPES pH 7.5, 200 mM NaCl, 5% glycerol). Protease was removed from the solution using Glutathione Sepharose 4 Fast Flow beads (Cytiva). After a polishing step using cation exchange HiTrap SP column (5 ml, Cytiva) the protein was dialysed back into buffer B. Concentrations were determined using Nanodrop One (Thermo Fischer Scientific). Folding state of the proteins was confirmed using circular dichroism (CD) spectra. Aliquots were snap-frozen in liquid nitrogen and stored at - 80°C. HHoF and HHoF histones from LC_3 were expressed and purified according to the same protocol as for HHoB histone. HMfB histone (*M. fervidus*) was expressed in *E. coli* LOBSTR cells by induction (0.5 mM IPTG at OD_600_ 0.5-0.6) and incubated at 37°C for 3-4 hours. The purification steps were identical to HHoB, with an additional heat incubation step of the supernatant (after lysis) at 80°C for 15 mins in a waterbath. This helped enriching for the thermostable HMfB histone.

#### DNA sequences and preparation

The Widom601^1^ derived 147 bp sequence was used for EMSAs and cryoEM: CTGGAGAATCCCGGTGCCGAGGCCGCTCAATTGGTCGTAGACAGCTCTAGCACCGCTTAAACGCACGTACGCGCTGTCCCCCGCGTTTTAACCGCCAAGGGGATTACTCCCTAGTCTCCAGGCACGTGTCAGATATATACATCCTGT.

A short DNA of random sequence and length 33 bp DNA with 6-FAM on the 3’ end: AGGGTCACATGGGTGTTTGGCACTACCGACAGT-6-FAM was used for Micro-scale thermophoresis (MST). The oligos for the 33 bp were ordered from Sigma (HPLC-purified), and annealed together by heating up to 98°C, and then were gradually cooled down.

Native LC_3 sequence (420 bp length) from gene HeimC3_31310 was used for EMSA and cryo-EM: ACCAGTTTTATTAGATCAACATATCAAGAAGTTACTAAAATCCGTTAAAAAAATAACAACACCCTGAATTTGACCGTAGTGATTAAGCTCAGTAATGGAATAAAGGAGAAAAAAATTTTGAAATTTAAGAAGAAGATTTATTTTTGAAACAGCGACTACGAAGAAAAAAATTTAAAAGTCAAGCTATTTTATTTGGAAGCAGACAAAGTAACGTCTGTTTCTTTTATTGTTCGACGACCGCTGTTTCGTGCAATTTCTACTGAATATCTGGCTACTTCGAGACCACGCTCTCCTAATAAATCGTTCAGTGCTTTAATTGCACCACTTGATACACGGAAAGCGCCGGCTTCACGGATTAATTTCTCTACTTTTGCAGAAGCGAAAGCTTGTCCTTTAGATCGGCTAGATCTACGTCGACTT

DNA constructs were ordered from GeneArt as parts of pMA or pMK vectors. DNA templates of interest were amplified by PCR, and purified via ion exchange (Resource Q column 5 ml, Cytiva). Force spectroscopy measurements were performed using biotinylated double stranded λ DNA of length 48.5 kb (SKU 00001, Lumicks).

#### In-vitro reconstitution and binding affinity measurements

Histone-DNA complexes were reconstituted in reaction buffer containing 20 mM HEPES pH 7.5, 100 mM NaCl, 5% glycerol. DNA concentration was kept constant (18 nM) and protein concentration was varied (9 – 540 nM). The reaction was incubated for 20 min at RT, then put on ice, and samples were loaded into the wells of a 5.5% 0.5x TBE PAGE gel. EMSAs were run at 80V for 90 min in 0.5x TBE buffer at 4°C. EMSA gels were then stained with SYBR gold (Invitrogen) and imaged in a Typhoon FLA 9500 (GE) imager. To study histone-DNA binding in the presence of magnesium ions, the reaction buffer and running buffer were supplemented with 10 mM MgCl2 and EMSAs were run for 120 min at 80 V.

EMSA with DNA substrates of different lengths was done by mixing GeneRuler Ultra Low Range Ladder with HHoB at the indicated w/w ratios in 20 mM HEPES pH 7.5, 100 mM NaCl, 5% glycerol (with 0 or 10 mM MgCl2). Samples were incubated at RT for 30 minutes before being run in 10% TBE-polyacrylamide gel at 120 V for 60 minutes (105 minutes for samples with MgCl2) at 4 C.

The binding affinity of HHoB was quantified using MST with a random 33 bp DNA oligonucleotide as the substrate. Experiments were conducted using a Monolith NT.115 (NanoTemper Technologies) with "blue" excitation at 40% LED power. The DNA substrate was used at a final concentration of 15.5 nM. Samples were prepared in a buffer containing 20 mM HEPES (pH 7.5), 100 mM NaCl, and 0.05% Tween-20, with an additional condition in which the buffer was supplemented with 1 mM MgCl₂. Measurements were performed in triplicates using Monolith NT.115 MST Premium Coated Capillaries (K005, NanoTemper Technologies). Data were analyzed using the Kd fit model in MO.Affinity analysis software (NanoTemper Technologies).

#### Sample preparation for cryo-EM

Nucleosomes were reconstituted by mixing DNA and histone proteins at a 1:20 molar ratio (DNA concentration 1.9 µM) in buffer A containing 20 mM HEPES (pH 7.5) and 100 mM NaCl. For magnesium concentration screening, nucleosome reconstitution was performed in buffer A supplemented with 1, 20, 40, 60, 80, or 100 mM MgCl₂. Samples were incubated at room temperature (RT) for 20 min and subsequently placed on ice until plunge-freezing. Cryo-EM grid preparation was performed using Quantifoil R 2/1 Cu 200 mesh grids, which were glow-discharged for 20 s at 0.26 mbar pressure and 25 mA current using a PELCO easiGlow (Ted Pella) device. A 3 µl aliquot of the sample was applied to the grids inside a Vitrobot Mark IV (Thermo Fisher Scientific) at 100% humidity and 20°C. Excess liquid was blotted away with a blot force of 7 for 2.5 s, and the grids were vitrified by plunging into liquid ethane.

#### Data collection

Data collection of HHoB – DNA complex in buffer A was performed on a 300 kV Titan Krios microscope (FEI) with a K2 summit direct electron detector (Gatan) in counting mode. A quantum energy filter (Gatan) was used with a slit width set to 20 eV. 3660 movies were collected with Serial EM software^2^, with defocus ranging from −1.5 to -2.5 μm at a nominal magnification of 130 kx and a pixel size of 1.04 Å. A constant stage tilt of 25° was applied during acquisition in order to compensate for “preferred” orientation of the particles. The total electron dose of 46.63 e^−^/Å^2^ was distributed over 40 frames. At the same microscope, datasets were also collected for the magnesium screen – these involved micrographs for samples in 20 mM, 40 mM, 60 mM and 80 mM MgCl2, taken with the same acquisition parameters of 65.04 e^−^/Å^2^ dose, 130kx magnification and defocus range of −0.5 to −1.75 μm. Data Collection for HHoB mut1 and HHoB mut2 in 1 mM and 100 mM MgCl2 respectively were also carried out on the same microscope. HHoB mut1 dataset was collected at 130 kx magnification and pixel size of 1.04 Å. A dose of 62.44 e ^−^/Å^2^ was spread over 40 frames. HHoB mut2 dataset was collected at 105 kx magnification and pixel size of 1.33 Å. A dose of 51.87 e ^−^/Å^2^ and this was distributed over 35 frames. The defocus range in both these datasets was −0.5 to −1.75 μm.

Data collection of HHoB – DNA complex in buffer containing 1 mM MgCl2 was performed on a G4 Titan Krios microscope (FEI) equipped with a Falcon4i direct electron detector (Thermo Fischer Scientific) and Selectris X Energy filter. Data were collected with Serial EM software^2^, with defocus ranging from −0.5 to −1.75 μm at a nominal magnification of 165 kx and a pixel size of 0.73 Å. Energy filter slit width was set to 20 eV. The total electron dose of 59.44 e^−^/Å^2^ was distributed over 40 movie frames recorded in EER format.

Micrographs of HHoB – DNA complex in buffer A supplemented with 20 mM MgCl2 were recorded on a 300 kV Titan Krios microscope with a K3 detector and Quantum energy filter. Data were collected with SerialEM software, with a defocus range of −0.5 to −1.5 μm at 105 kx magnification and pixel size of 0.82 Å. The width of the energy filter was set to 20 eV with a total electron dose of 63.84 e^−^/Å^2^. This was distributed over 40 frames. Micrographs for processing HHoB in 100 mM MgCl2 were collected on the same microscope, with a defocus range of –0.5 to –1.5 μm at 130 kx magnification and pixel size of 0.645 Å. The width of the energy filter was set to 20 eV with a total electron dose of 39.93 e^−^/Å^2^. This dose was distributed over 40 frames.

#### Data processing and analysis

Data processing for all datasets was carried out in cryoSPARC^3^ (v.4.4.1).

*Dataset 1* (buffer A – no MgCl2): After preprocessing, 4,300 particle picks were curated from a blob picker by 2D classification and Topaz^4^ model was trained using these particles – they were used for picking particles from the entire dataset and another topaz model was trained on these picks after 2D classification. 318,063 particles were then used for 2D classification to eliminate junk particles, and followed by ab-initio reconstruction, and 3D classification into 3 classes. One well-resolved class was then selected, and it contained 96,361 particles. The map was processed by non-uniform refinement and per-particle CTF correction to yield the final map at 4.4 Å. 3DFSC server^5^ was used to ensure isotropic resolution of the structures.

*Dataset 2* (buffer A + 1 mM MgCl2): After preprocessing, 50k particles were curated after blob and template picking. The dataset showed preferred orientation for top views, and large heterogeneity in conformations, so two topaz models were trained – one for top views and another for side and oblique views. In total, 3,108,514 particles were picked and filtered through two rounds of 2D classifications into open and closed conformations. Then ab-initio reconstruction and 3D classification (into 2 classes each) for open and closed conformations were performed separately. Good classes (96,738 particles for open and 117,260 particles for closed conformation) were processed by non-uniform refinement and post processed by CTF and reference-based motion correction to yield the final maps at 3.6 Å and 3.5 Å for the open and closed conformation respectively. Additionally, 546,206 particles were aligned in 3D by homogenous refinement to a low-resolution mixed conformation. This was subjected to 3D classification to obtain low resolution maps of a range of mixed conformations – highlighting the heterogeneity of the dataset (Figure S4).

*Dataset 3* (buffer A + 20 mM MgCl2): After preprocessing, 7,816 particle picks were curated from a blob picker by 2D classification and Topaz model was trained using these particles – they were used for picking particles from the entire dataset and another topaz model was trained on these picks after 2D classification. 214,840 particles were then used for 2D classification to eliminate junk particles, followed by ab-initio reconstruction and 3D classification, which resulted in one well-resolved class containing 40,918 particles. From the micrographs and 2D classes it can be appreciated that the dataset indeed contained open hypernucleosomes that are very flexible and contain mixed conformations. The map was processed by non-uniform refinement and per-particle CTF corrections to yield the final map at 10.5 Å.

*Dataset 4* (buffer A + 100 mM MgCl2): After pre-processing, 4,964,843 particles were picked using filament tracer. After three rounds of 2D classification 484,756 particles were selected for ab initio volume generation and refinements. Initially a cylinder with outer and inner diameter 13 nm and 1 nm respectively was used as template for refinement without helical parameters. Symmetry search was performed and used for helical refinement. The particles were then 3D classified and the well-resolved class (353,468 particles) was helically refined with non-uniform refinement enabled. After per group CTF refinement and reference-based motion correction, final helical refinement was carried out to obtain the HHoB closed hypernucleosome map at 2.6 Å. Helical twist and pitch was determined to be 77.96° and 19.42 Å respectively.

*Dataset 5* (magnesium concentration screen): Thirty micrographs from the 40 mM MgCl₂ dataset were pre-processed and evenly divided into two subsets. Each subset was used to train a separate Topaz model following blob-picking and 2D classification. These independently trained models were then applied to pick particles from 30 micrographs collected at 20, 40, 60, and 80 mM MgCl₂, as well as from 30 micrographs randomly selected from the 0, 1, and 100 mM MgCl₂ larger datasets. We note, that the 1 mM dataset was collected at a 0.73 Å pixel size, the 100 mM at 0.64 Å pixel size, and the rest (0, 20, 40, 60, 80) at 1.04 Å pixel sizes. The particle box extraction was adjusted accordingly. Following particle picking, all particles underwent 2D classification to remove junk particles and enhance data quality. After this filtering step, the number of particles per dataset ranged from 1,349 to 7,575. Particles were categorized into open, mixed, or closed conformations based on their 2D class averages. Representative 3D reconstructions of these conformational states, derived from the larger 1 mM MgCl₂ dataset (dataset 2), are presented in Figure S4. Fast Fourier Transform (FFT) analyses of the micrographs (shown in Figures S5 and S10) were performed using FFT function in FIJI software^6^ (version 2.16.0).

#### Model fitting and refinement

Initially, the AlphaFold2 prediction of the histone HHoB dimer was used for rigid body fitting in UCSF ChimeraX^7^ v.1.7.1. After real space refinement in Phenix^8^ (v1.21.1), per residue fitting was done in Coot ^9^ v 0.9.8.93 EL using all-molecule self restraints and finally the model was relaxed into the density using ISOLDE^10^ in ChimeraX. For the open conformation model, DNA was fit into the densities by ISOLDE relaxation from H3-H4 open eukaryotic octasome structure (PDB 7x58^11^) with DNA H-bond restraints. Nucleic acid backbone angles were refined with DNA B-form restraints in Coot. RMSD and surface area calculations were performed in ChimeraX.

#### Protein labelling with fluorescent dye

To label HHoB mut3, it was incubated at molar ratio 1:5 with Atto647N-maleimide (FP-202- 647N, Jena Biosciences) overnight at RT on a shaker in buffer A. Free dye was removed from buffer using desalting column (PD-10, Cytiva). The degree of labeling was estimated with the Nanodrop (Thermo Fisher Scientific) to be 0.99 (99 %). Aliquots were snap frozen in liquid nitrogen and stored at -80℃. DNA binding properties of labelled HHoB were the same as for unlabelled according to EMSA (not shown).

#### Force Spectroscopy measurement and analysis

Single-molecule force-extension measurements were performed using a high-resolution correlative fluorescence optical tweezers instrument (C-trap, LUMICKS). A microfluidic flow cell (LUMICKS) with five parallel laminar flow channels was used, allowing the controlled movement of the optical traps between different solutions. Streptavidin-coated polystyrene beads (4.35 µm in diameter) were trapped in phosphate-buffered saline (1xPBS) and used to anchor biotinylated double-stranded λ DNA (48.5 kb) for force measurements. The traps were calibrated to have a stiffness of 0.3 to 0.4 pN/nm. The first baseline measurement was taken prior to DNA anchoring in the buffer channel containing Buffer C (20 mM HEPES pH 7.5, 100 mM NaCl, 0.1% BSA). Force-extension measurements were performed by stretching the DNA from a low-force regime (L= 8 µm) to a high-force, stretched state (L= 16.7 µm) at a pulling speed of 0.5 µm/s. Following this, the anchored DNA was incubated in the protein channel containing 200 nM histone in the same buffer for 5 minutes at an inter-bead distance of 8 µm. Force-extension measurements were then performed on the histone-DNA complex by stretching from the relaxed to stretched state at a pulling speed of 0.05 µm/s. After histone incubation and force-extension measurements, the DNA was ruptured by further bead separation to ensure complete removal, and a second baseline measurement was recorded in the protein-containing channel. The first baseline was used to correct force-extension data for free DNA, while the second was used to correct for the histone-DNA complex. All data were processed and analyzed in Python using the LUMICKS Pylake library (version 1.6.1). Statistical comparisons were performed using paired t-tests in JASP (version 0.19.3) to assess force differences between datasets.

#### Tethered Particle Motion

Tethered Particle Motion (TPM) experiments were done following the procedure described in^12^ on a 685 bp DNA substrate with a 47% GC content in 20 mM HEPES, pH 7.5, 200 mM NaCl, 5% glycerol. Each measurement was done in duplicate. To select for single-tethered beads, an anisotropic ratio cutoff of 1.3 and a standard deviation cut-off of 8% were used. Data analysis was done as described in Hu, et al. (2024). The end-to-end distance was calculated by subtracting the mean bead radius from the largest 5% XY-displacement of all beads in each measurement.

#### Multi-Sequence Alignment

The histone sequences of Hodarchaea, Odinarchaea, Lokiarchaea, Thorarchaea, and Euryarchaea were obtained from Uniprot^13^ and Clustal Omega^14^ was used for generating multi-sequence alignment. Their respective UniProt IDs are mentioned in Figure S12. Software JalView^15^ (v 2.11.4.1) was used for visualization.

---

**Figure S1.**
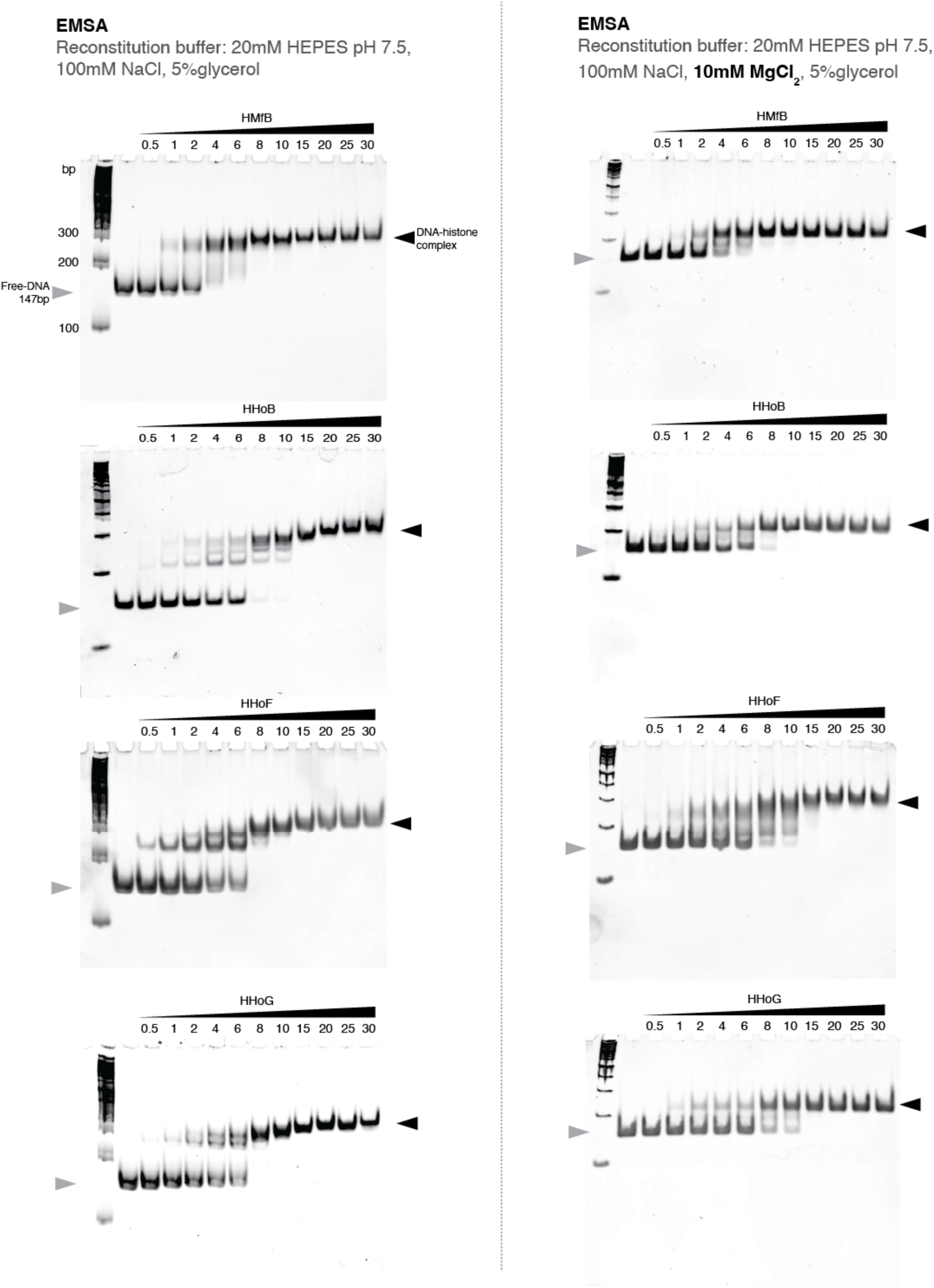
Related to Figure 1. Electrophoretic mobility shift assay (EMSA) gels showing in-vitro reconstitutions of 147 bp Widom601 DNA with different histones in buffer A (left) and buffer A supplemented with 10mM MgCl2 (right). Increasing molar ratios of histone to DNA are shown on top of the gels. DNA concentration was constant (20nM). First lane shows 100bp DNA ladder (peqGOLD 10Obp Plus).

**Figure S2.**
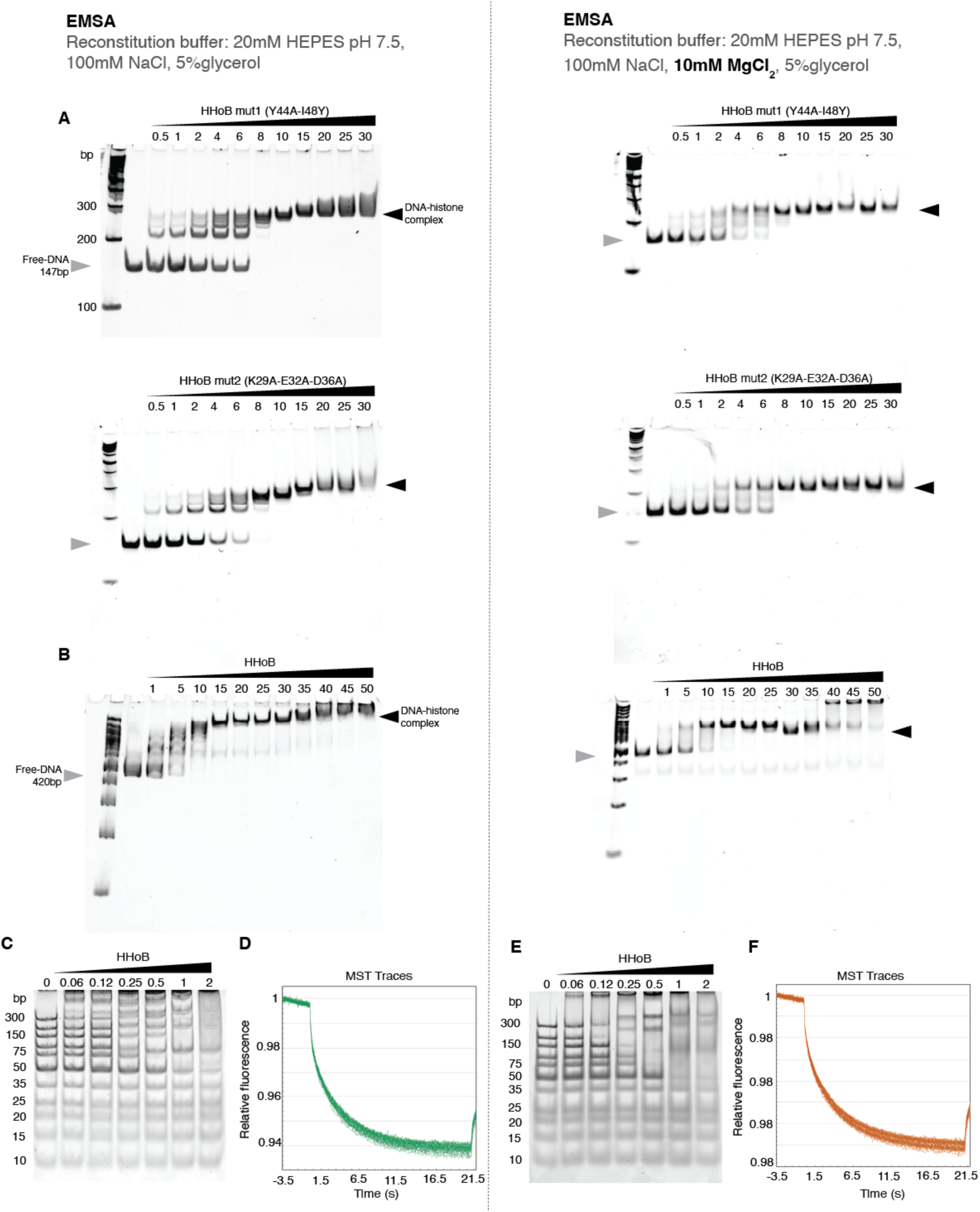
Related to Figure 1. EMSA gels showing in-vitro reconstitutions of (A) HHoB mutants with 147 bp Widom601 and (B) HHoB with 420bp HeimC3_31310 native DNA. Increasing molar ratios of histone to DNA are shown on top of the gels. DNA concentration was constant (20nM). First lane shows 100bp DNA ladder (peqGOLD 100bp Plus). (C,E) HHoB with GeneRuler Ultra Low Range ladder DNA with in buffer A (left) and buffer A supplemented with 10mM MgCl_2_ (right). (D, F) show MST traces of HHoB with 33bp 6-FAM labelled DNA in buffer A and buffer A supplemented with 1mM MgCl_2_.

**Figure S3.**
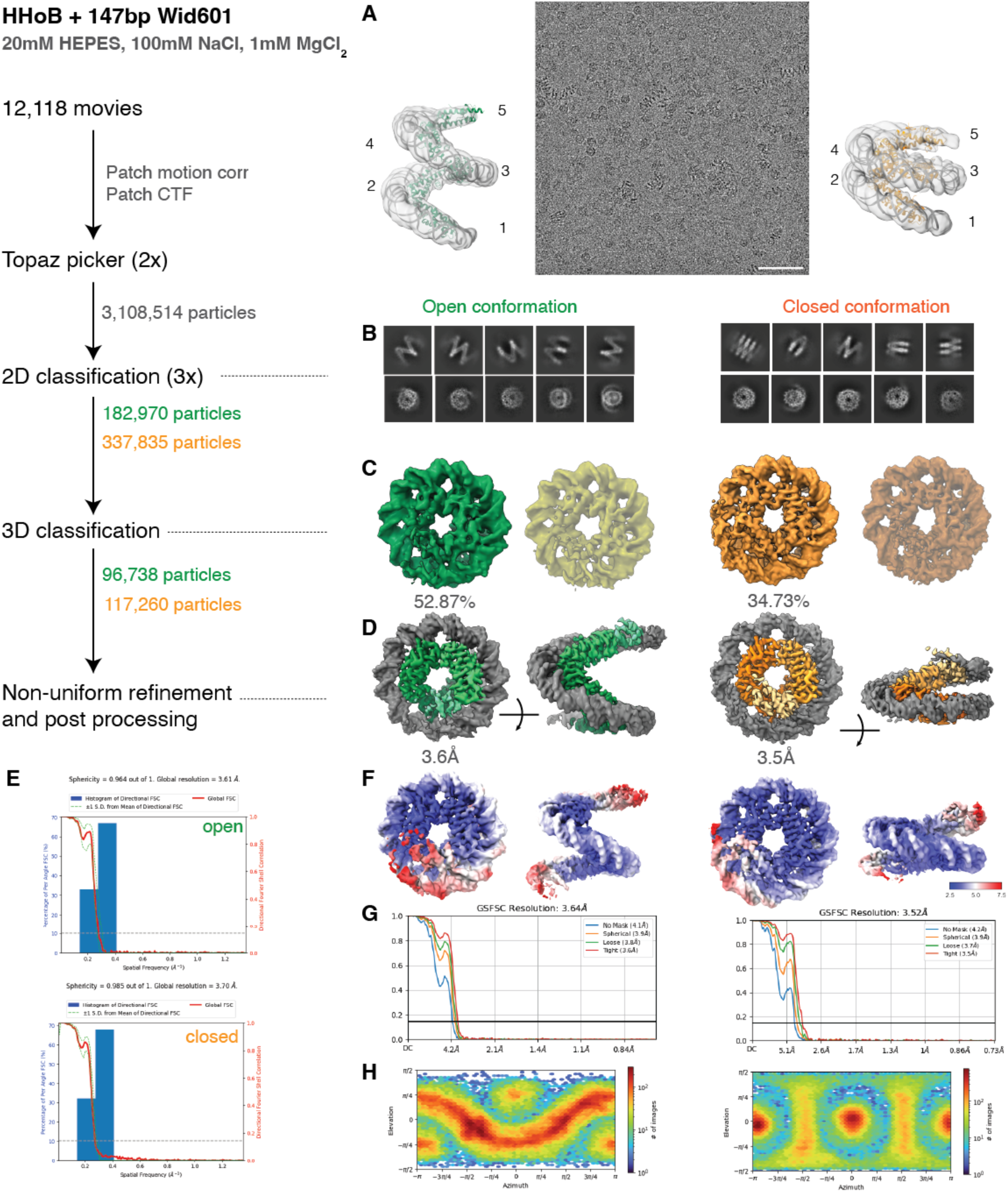
Related to Table 1. Single-particle Cryo-EM processing of the HHoB-DNA complex in 1 mM MgCl_2_. The flowchart illustrates the image processing workflow, with dotted lines connecting to corresponding figures for each step. (A) A representative raw micrograph with refined densities, gaussian-filtered and fitted with five histone dimers shown on the sides. Scale bar= 50 nm, (B) Representative 2D class averages displaying both open and closed conformations of the complex. (C) Results from 3D classification, with discarded classes rendered at lower opacity. (D) Refined densities visualized in two orthogonal views. (E) 3D Fourier shell correlation (FSC) curves demonstrating isotropic resolution distribution. (F) Local resolution maps indicating the variation in resolution across the reconstructed density. (G) Gold-standard FSC (GSFSC) curves with a cutoff at 0.143, providing an assessment of overall map resolution. (H) Particle angular distribution plots.

**Figure S4.**
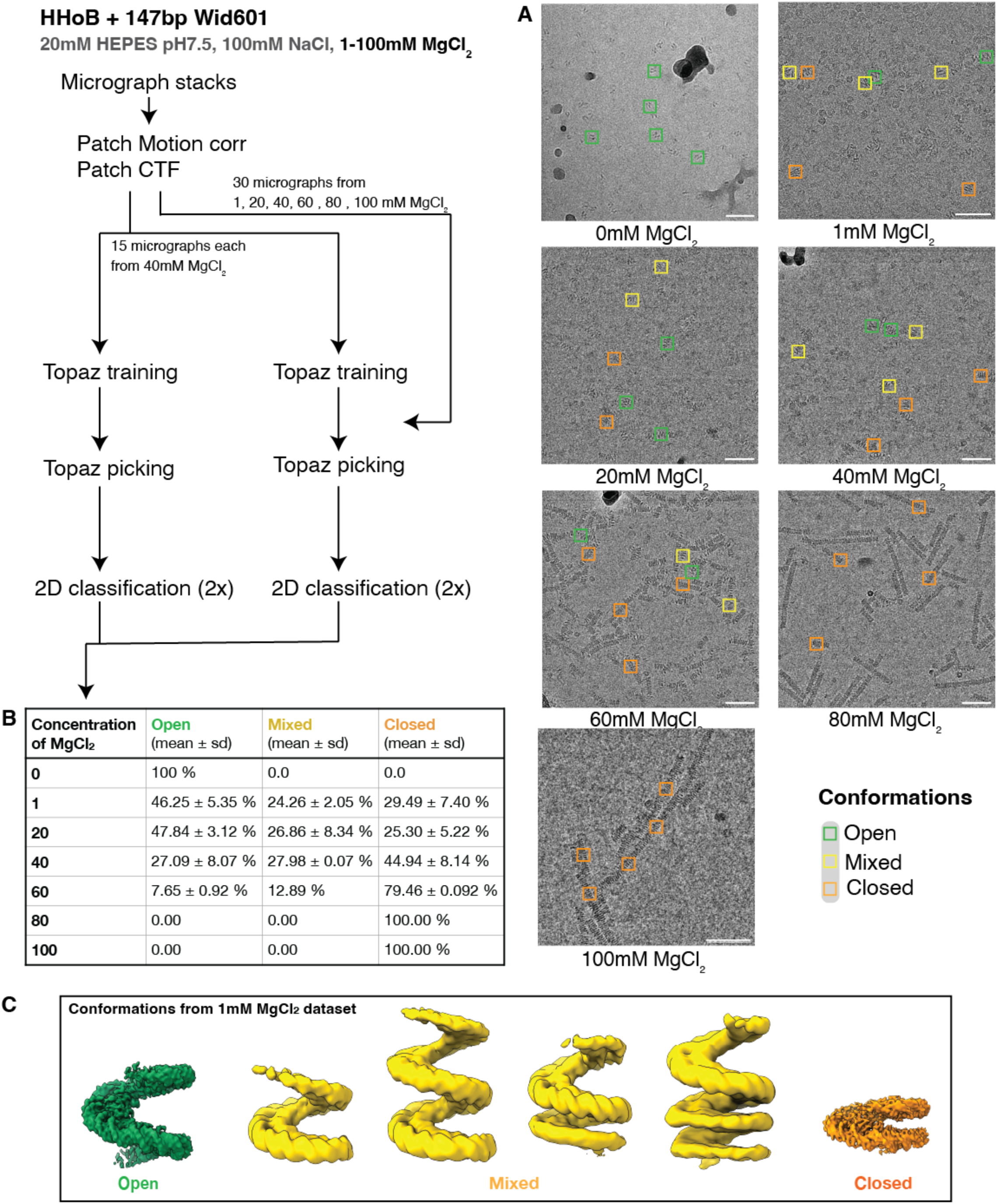
Related to Figure 2. Single-particle analysis of the HHoB-DNA complex in buffers with varying MgCl concentrations. The flowchart (left) outlines the image processing pipeline, while (A) representative micrographs display some selected particle picks annotated by their assigned conformational states. Scale bar = 50nm. ( Green = open, yellow = mixed and orange = closed conformations). (B) Summary table presenting the distribution of particles across these conformations, with values reported as the mean ± standard deviation, based on technical duplicates generated by two independently trained Topaz particle-picking models. (C) A gallery of conformations obtained from the 1mM MgCl2 dataset.

**Figure S5.**
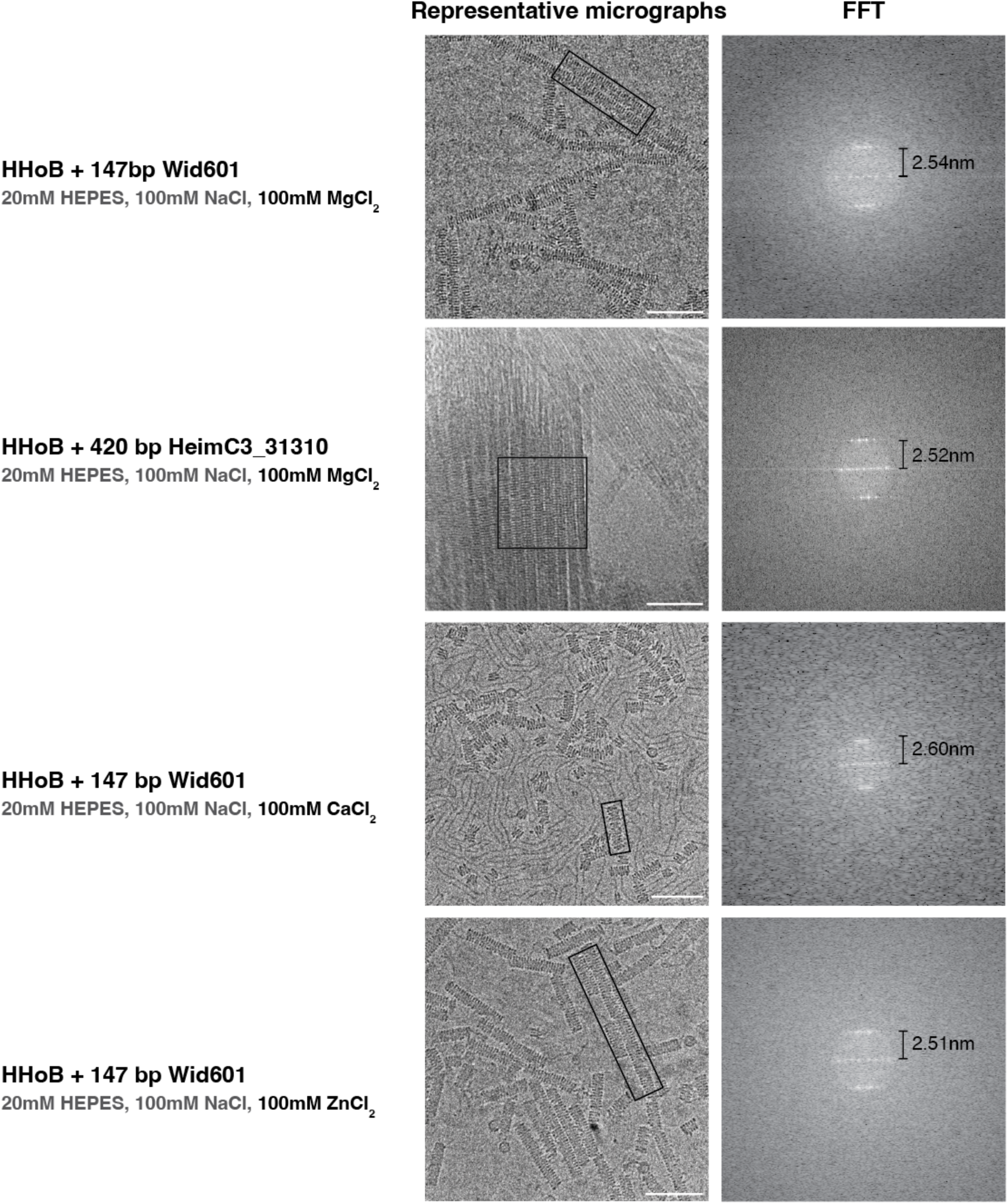
Related to Figure 2. HHoB closed hypernucleosomes formed in different conditions (indicated on the left). Representative micrographs display annotated regions of interest, highlighting the presence of closed hypernucleosomes under different buffer conditions. The right panel presents the corresponding fast Fourier transform (FFT) analysis, with inter-turn distances, "pitch", annotated. Scale bar is 50 nm.

**Figure S6.**
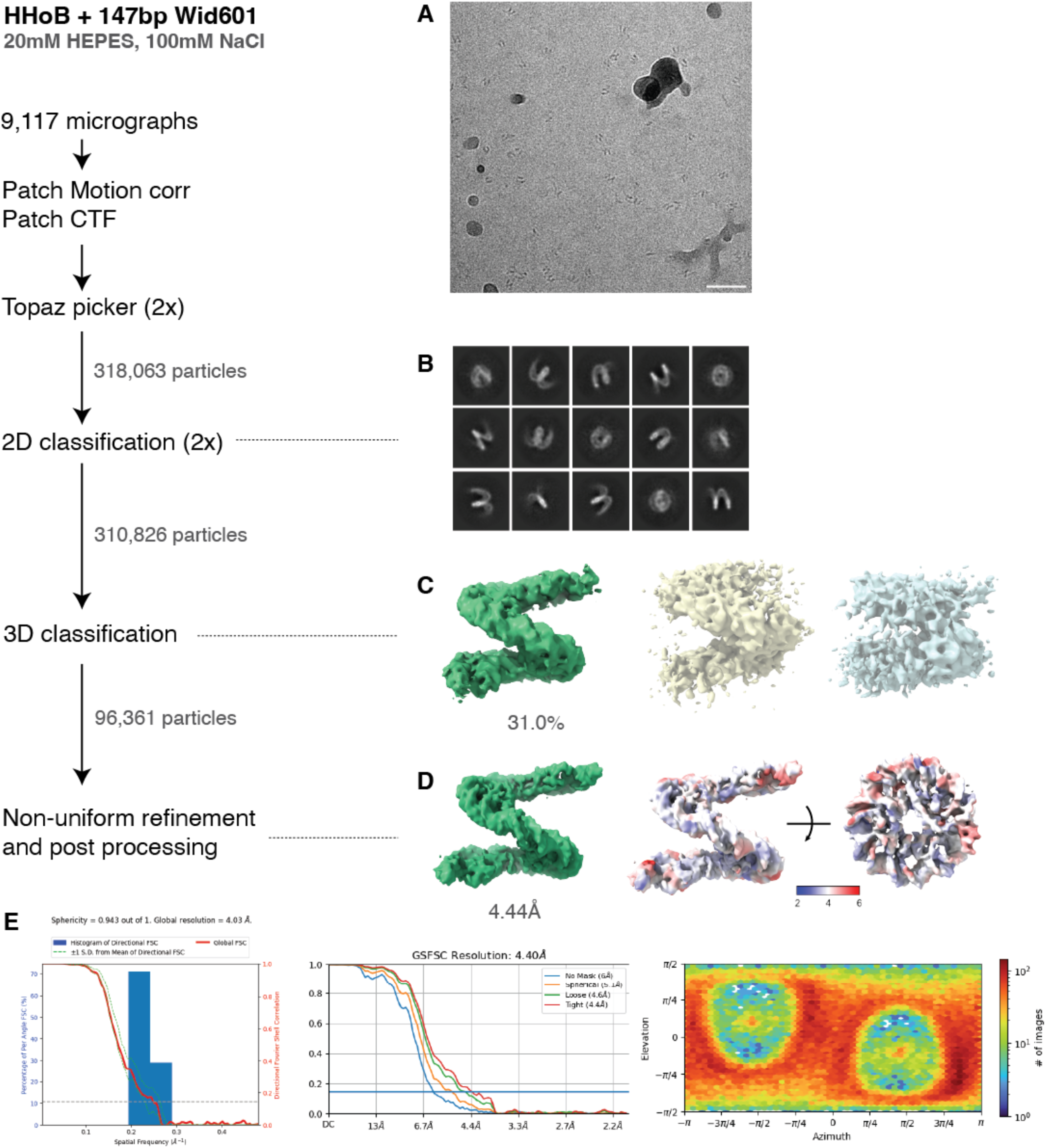
Related to Table 1. Single-particle Cryo-EM processing of the HHoB-DNA complex in buffer A. The flowchart illustrates the image processing workflow, with dotted lines connecting to corresponding figures for each step. (A) A representative raw micrograph. Scale bar: 50nm. (B) Representative 2D class averages, (C) Results from 3D classification, with discarded classes rendered at lower opacity. (D) Refined density along with local resolution map in two views. (E) On the left - 3DFSC curve demonstrating isotropic resolution distribution, middle - GSFSC curve with a cutoff at 0.143, providing an assessment of overall map resolution and right - particle angular distribution plot.

**Figure S7.**
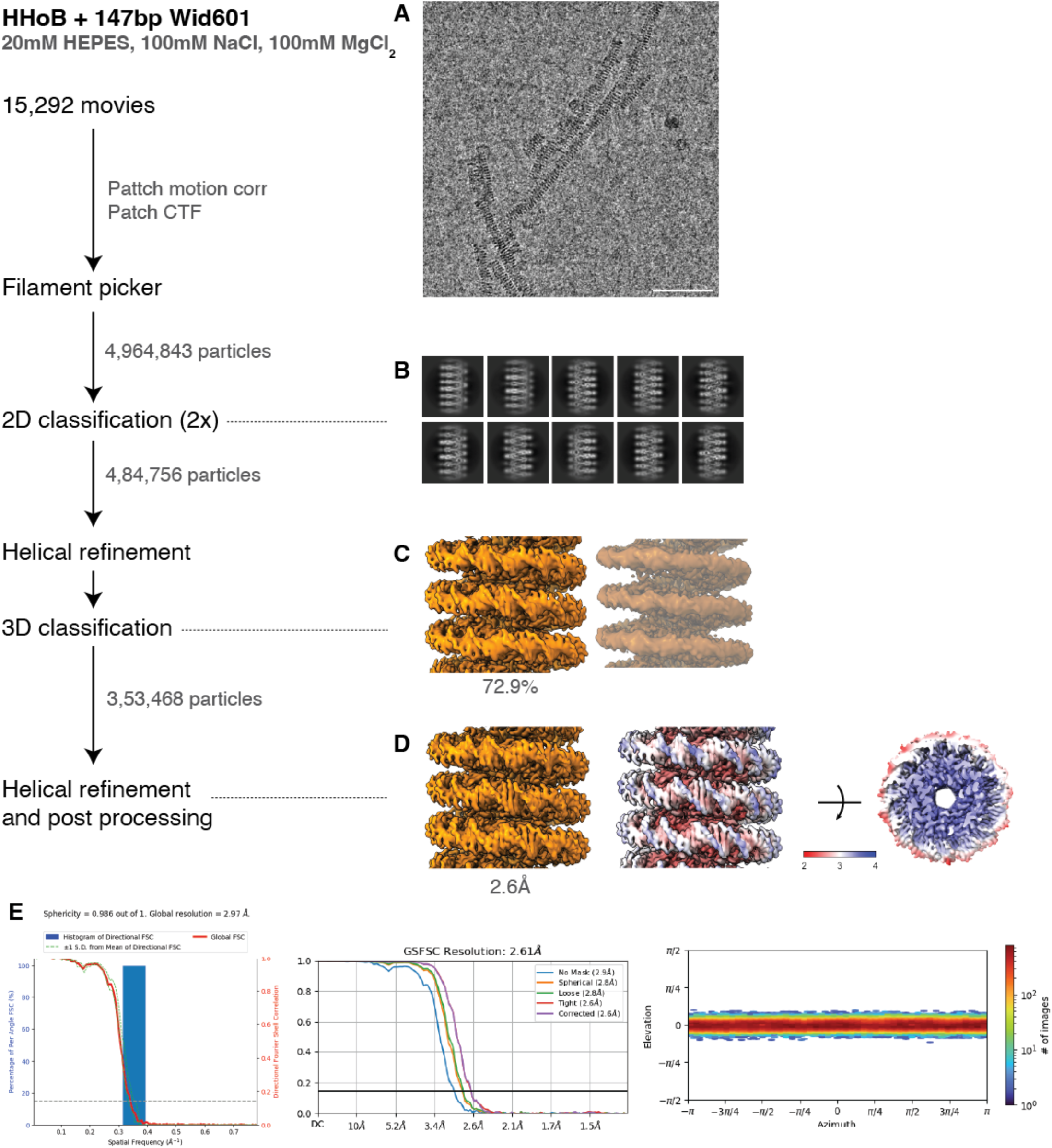
Related to Figure 3. Single-particle Cryo-EM processing of the HHoB-DNA complex in 100mM MgCl_2_. The flowchart illustrates the image processing workflow, with dotted lines connecting to corresponding figures for each step. (A) A representa­ tive raw micrograph. Scale bar: 50nm. (B) Representative 2D class averages, (C) Results from 3D classification, with discarded classes rendered at lower opacity. (D) Refined density along with local resolution map in two views. (E) On the left - 3DFSC curve demonstrating isotropic resolution distribution, middle - GSFSC curve with a cutoff at 0.143, providing an assessment of overall map resolution and right - particle angular distribution plot.

**Figure S8.**
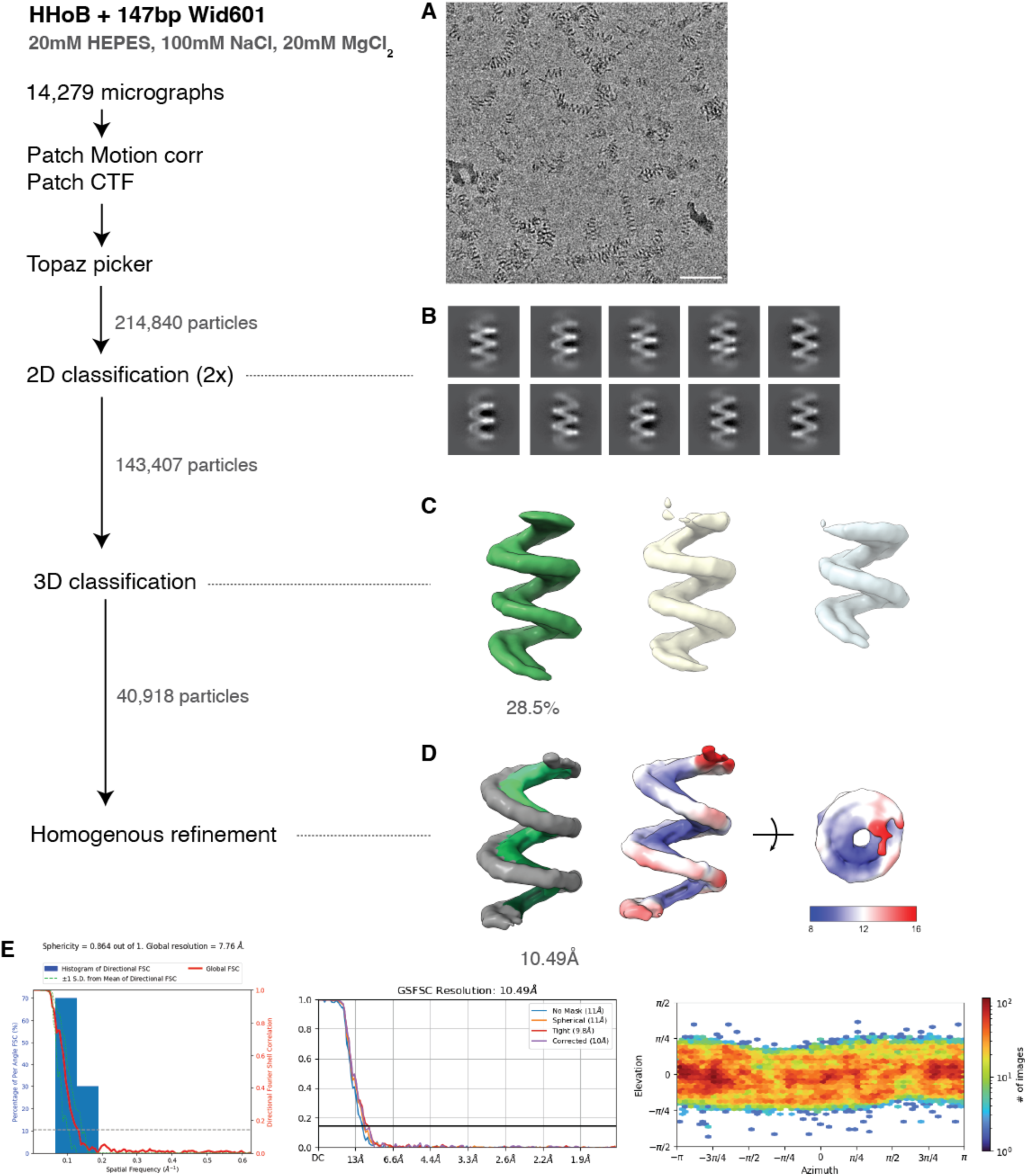
Related to Figure 3. Single-particle Cryo-EM processing of the HHoB-DNA complex in 20mM MgCl_2_. The flowchart illustrates the image processing workflow, with dotted lines connecting to corresponding figures for each step. (A) A representative raw micrograph. Scale Bar: 50nm. (B) Representative 2D class averages, (C) Results from 3D classification, with discarded classes rendered at lower opacity. (D) Refined density along with local resolution map in two views. (E) On the left - 3DFSC curve demonstrating isotropic resolution distribution, middle - GSFSC curve with a cutoff at 0.143, providing an assessment of overall map resolution and right - particle angular distribution plot.

**Figure S9.**
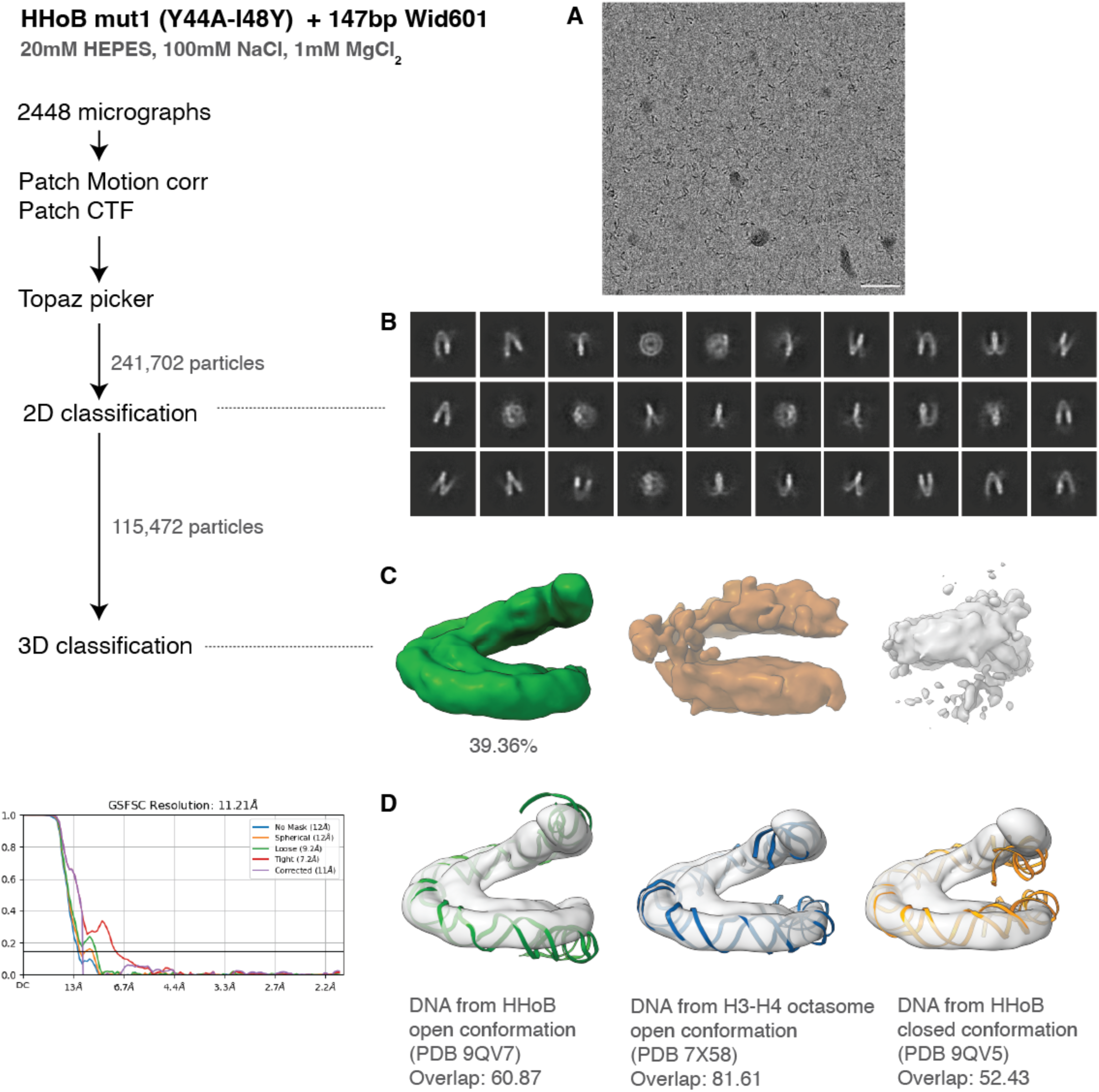
Related to Figure 5. Single-particle Cryo-EM processing of the HHoB mut1 - DNA complex in 1mM MgCl_2_. The flowchart illustrates the image processing workflow, with dotted lines connecting to corresponding figures for each step. (A) A representative raw micrograph. Scale bar: 50nm. (B) Representative 20 class averages, (C) Results from 30 classification, with discarded classes rendered at lower opacity. (D) Open conformation with rigid body fits of DNA from HHoB open model (green, PDB 9QV7), HHoB closed model (orange, PDB 9QV5) from this study, and H3-H4 octasome open model (blue, PDB 7X58). Overlap values were calculated in ChimeraX using fitmap.

**Figure S10.**
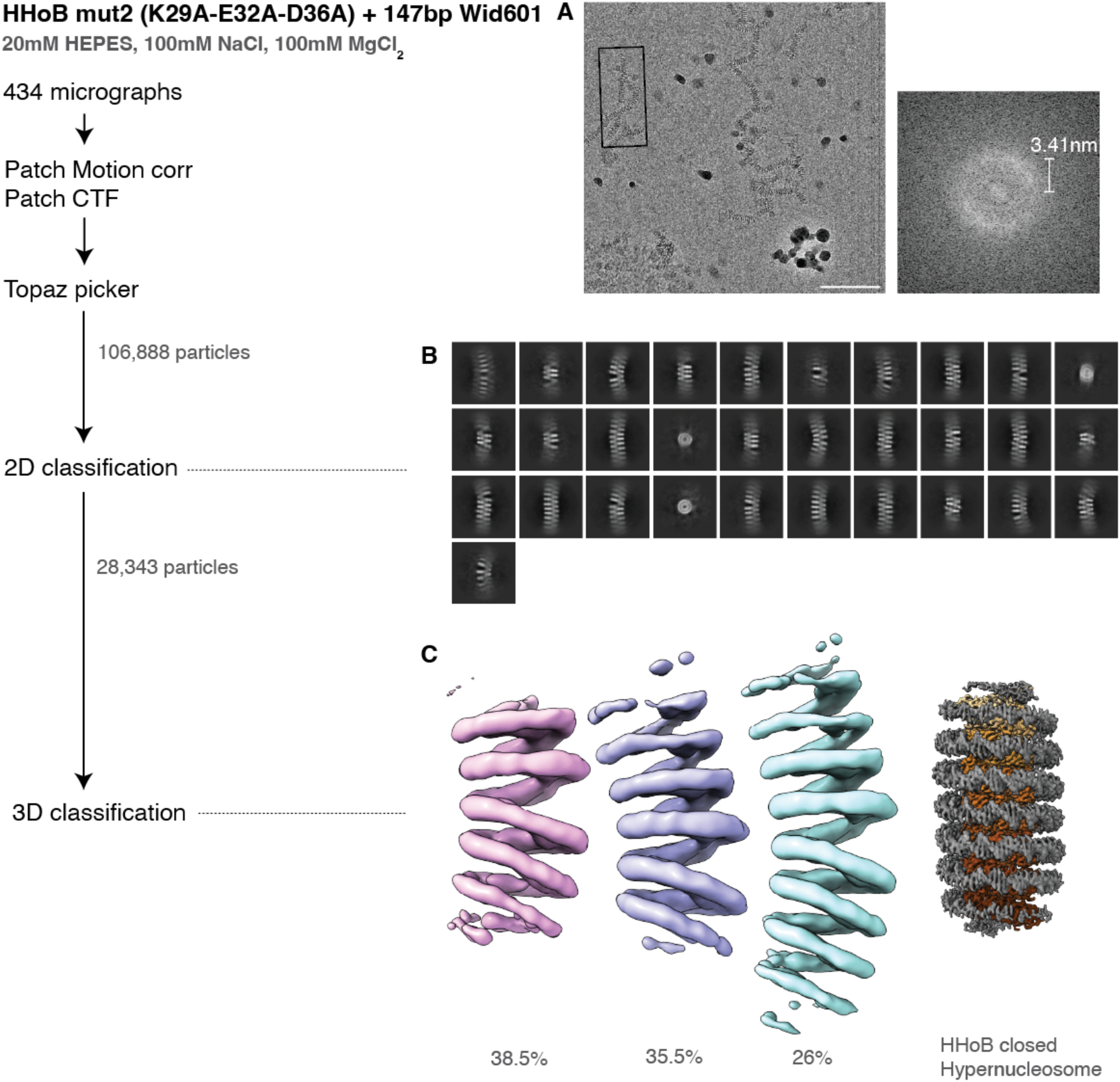
Related to Figure 5. Single-particle Cryo-EM processing of the HHoB mut2 - DNA complex in 100mM MgCl_2_. The flowchart illustrates the image processing workflow, with dotted lines connecting to corresponding figures for each step. (A) A representative raw micrograph.Scalebar= 50nm. The pitch of fibres in the box were determined by FFT (right). (B) Representative 2D class averages, (C) Results from 3D classification showing flexible fibres. HHoB closed hypernucleosome density is shown alongside for comparison.

**Figure S11.**
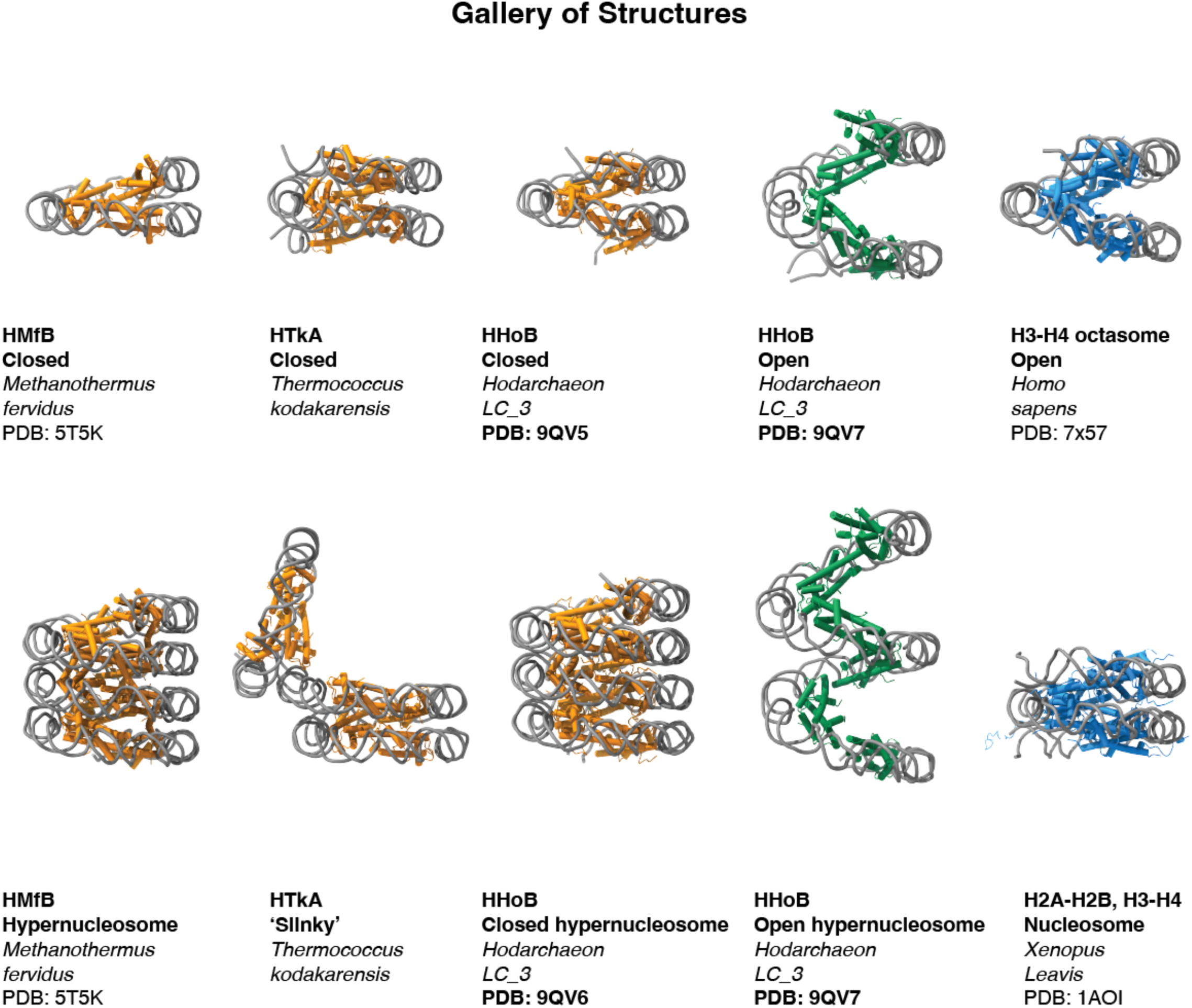
Related to Figure 6. Gallery of structures summarising known structural knowledge of histone-DNA complexes across archaea. From left to right: single unit and hypernucleosome inferred from crystal contacts of the histone HMfB (PDB 5T5K) from *M. fervidus.* Next, cryo-EM structures of the histone HTkA with DNA as published in ^10^ showing a canonical closed form and ’slinky’ conformations. The cryo-EM structures of the HHoB-DNA complex from this study are shown in closed (PDB 9QV5) and open conformations (PDB 9QV7), along with hypernucleosome fibres in the open (PDB 9QV7) and closed (PDB 9QV6) conformations. Lastly, the cryo-EM structure of H3-H4 octasome in open conformation (PDB 7X57) and the canonical eukaryotic nucleosome (PDB 1AOI) are shown in blue.

**Figure S12.**
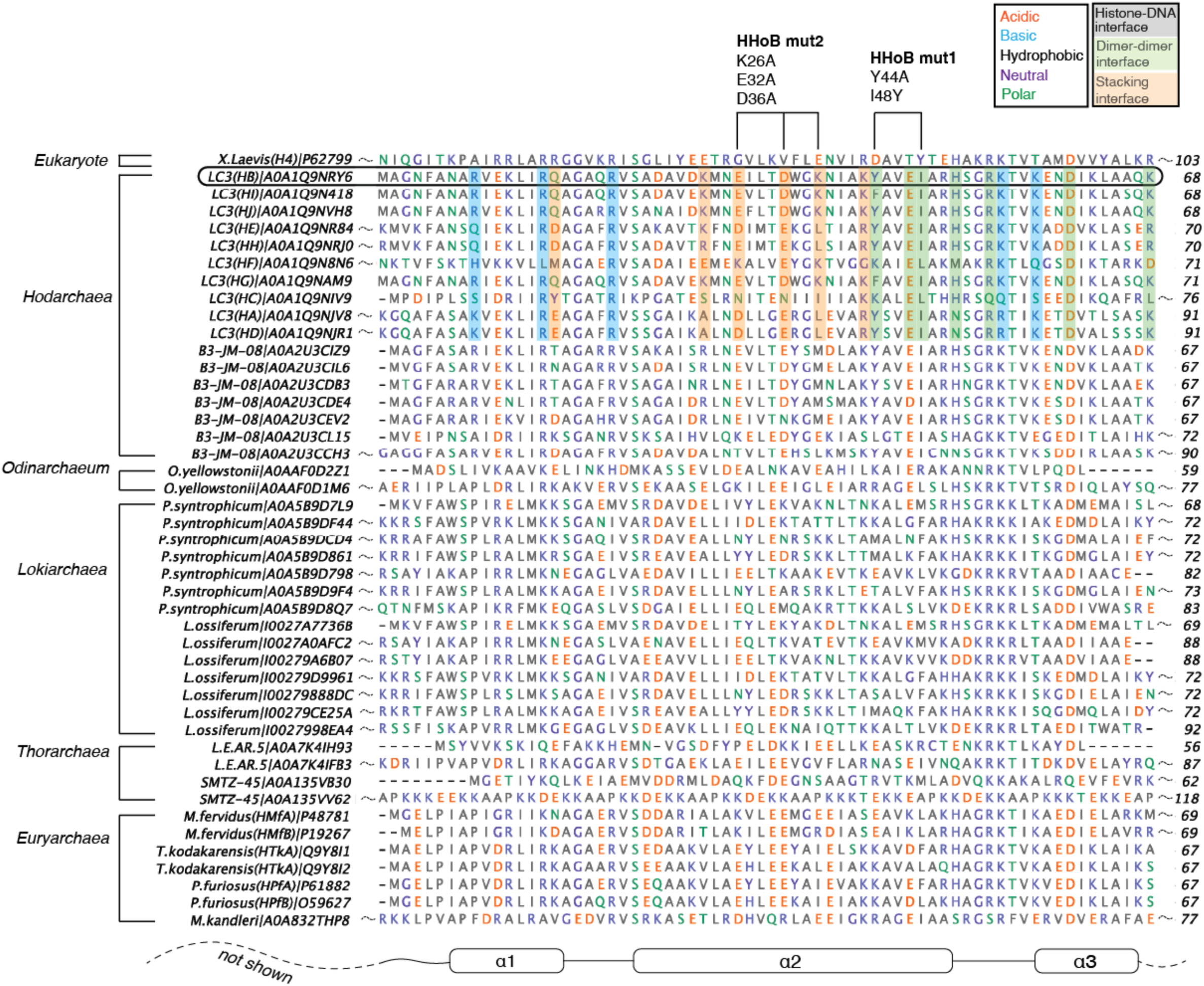
Related to Figure 6. Multi sequence alignment (MSA) of histone fold domain sequences across archaea - amino acids are colored by their physical properties (legend on the right) - and the histone HHoB is highlighted in the rounded box. Vertical shaded columns represent histone-DNA interactions (grey), dimer-dimer interactions (pale green) and stacking interactions (pale orange) for the *Hodarchaeon LC3* based on the HHoB nucleosome structures in this study.

**Supplementary Video S1:** Movie showing EM densities and associated models in this study. The names of the complexes and their buffer of reconstitution are mentioned on the bottom left, while the conformation of the complexes are denoted on the top right.

**Supplementary Video S2:** Movie showing the open (PDB 9QV7) and closed (PDB 9QV5) models and a morph between them at two histone dimers.

